# Image-Derived 3D Blood–Brain Mechanics: Cerebral Haemodynamics, Brain Motion and In Vivo Benchmarking

**DOI:** 10.64898/2026.08.24.746773

**Authors:** Yi Yang, Meiyao Wang, Yumin Liu, Wenbo Zhan, Daniele Dini, Tian Yuan

## Abstract

Cerebrovascular pulsatility drives measurable brain tissue deformation and has been associated with ageing and a range of neurological disorders. Yet how pulsatile haemodynamic forces are transmitted through deformable cerebral arteries into the surrounding brain remains poorly understood, particularly in anatomically realistic vascular geometries. Existing computational approaches have largely treated cerebral fluid and tissue mechanics separately or relied on idealised geometries, limiting our ability to determine how vascular anatomy simultaneously governs intraluminal haemodynamics and extravascular mechanical loading. Here, we develop an image-derived three-dimensional computational framework that jointly resolves pulsatile blood flow, arterial wall deformation and surrounding brain tissue motion in representative cerebral arteries. Four arterial segments, including the middle cerebral artery, middle cerebral artery bifurcation, basilar artery and internal carotid artery, are reconstructed from high-field (5 Tesla) magnetic resonance imaging data of a healthy subject. A finite-deformation fluid–structure interaction model is established by coupling non-Newtonian blood flow, hyperelastic arterial wall and hyper-viscoelastic brain tissue. The predicted tissue response is benchmarked against *in vivo* magnetic resonance elastography measurements of cardiac-induced volumetric strain over a cardiac cycle.

Results reveal spatially localised arterial and tissue deformation whose magnitude and distribution are strongly governed by vascular geometry and wall thickness. Among the segments examined, the internal carotid artery exhibits the largest deformation response, while reduced wall thickness increases strain transmission into the surrounding tissue. Geometrically complex regions also exhibit greater spatial heterogeneity in near-wall haemodynamic metrics. These findings demonstrate that cerebral vascular anatomy simultaneously shapes intraluminal haemodynamics and extravascular mechanical loading. By integrating image-derived vascular anatomy, coupled blood–vessel–brain mechanics and *in vivo* benchmarking within a unified framework, this study provides a mechanically consistent reference for healthy cerebral pulsatility and establishes a foundation for quantifying how blood–vessel–brain interactions are altered under pathological conditions.

**Highlights:**

- An image–derived framework is developed to quantify coupled blood–vessel–brain dynamics.
- Vessel geometry is shown to regulate local deformation and haemodynamics.
- Wall thickness is identified as a key determinant of pulsatile strain.
- A healthy benchmark is established for future cerebral disease modelling.

**Graphical Abstract:** 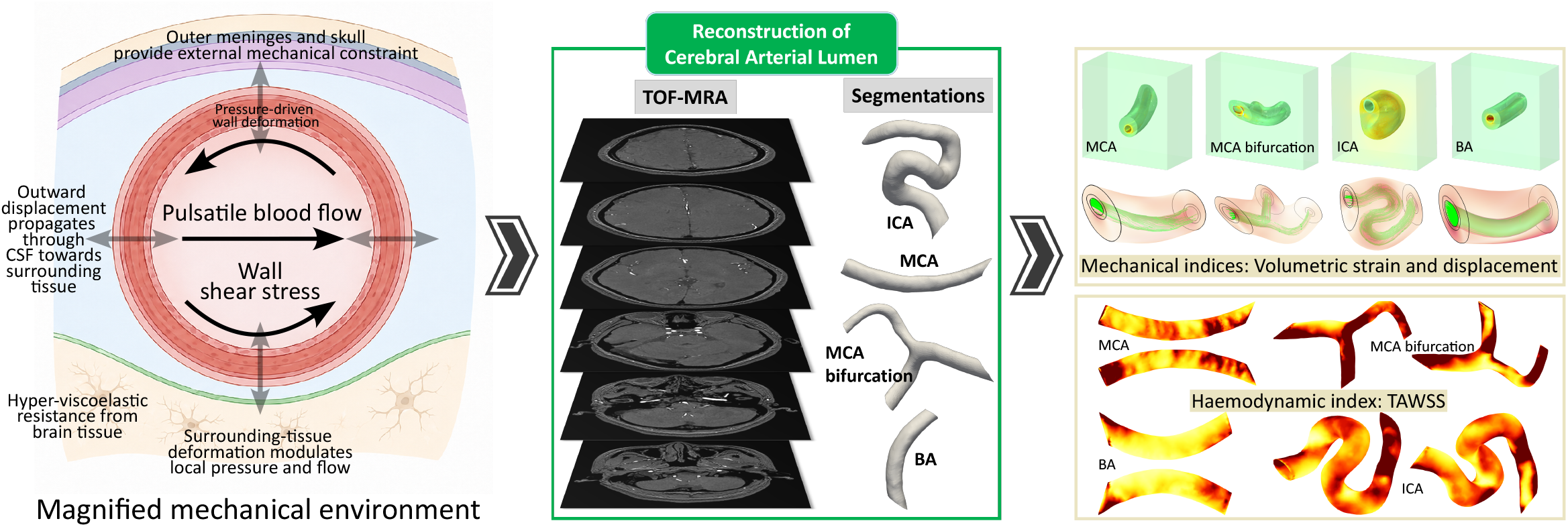

## 1. Introduction

Blood flow in the cerebral arterial system is subjected to continuous pulsatile loading driven by cyclic variations in blood pressure over the cardiac cycle [1, 2]. The resulting local haemodynamic environment, including pressure, velocity patterns and wall shear stress (WSS), is increasingly used to interpret vascular mechanobiology and disease-relevant remodelling pathways associated with conditions such as arterial stenosis, brain aneurysm formation and rupture risk [3, 4, 5]. Figure 1A illustrates the circle of Willis and the major cerebral arterial branches forming the anterior and posterior circulations. Unlike arteries in many other vascular beds, however, cerebral arteries operate within a highly constrained intracranial mechanical environment. As illustrated in Figure 1B, cerebral arteries are surrounded by cerebrospinal fluid (CSF) within the subarachnoid space and mechanically interact with the pia mater, adjacent brain parenchyma, meninges and, ultimately, the rigid skull. Pulsatile intraluminal pressure therefore not only drives blood flow but also deforms the compliant arterial wall and perturbs the surrounding intracranial environment. The resulting vascular motion is constrained by the surrounding fluid and tissue compartments, while deformation of these compartments can, in turn, modify the local mechanical conditions experienced by the vessel and flow, as illustrated in Figure 1C [6, 7, 8, 9]. Cerebral haemodynamics and extravascular tissue mechanics are therefore intrinsically coupled rather than independent phenomena [1, 10].

**Figure 1:**
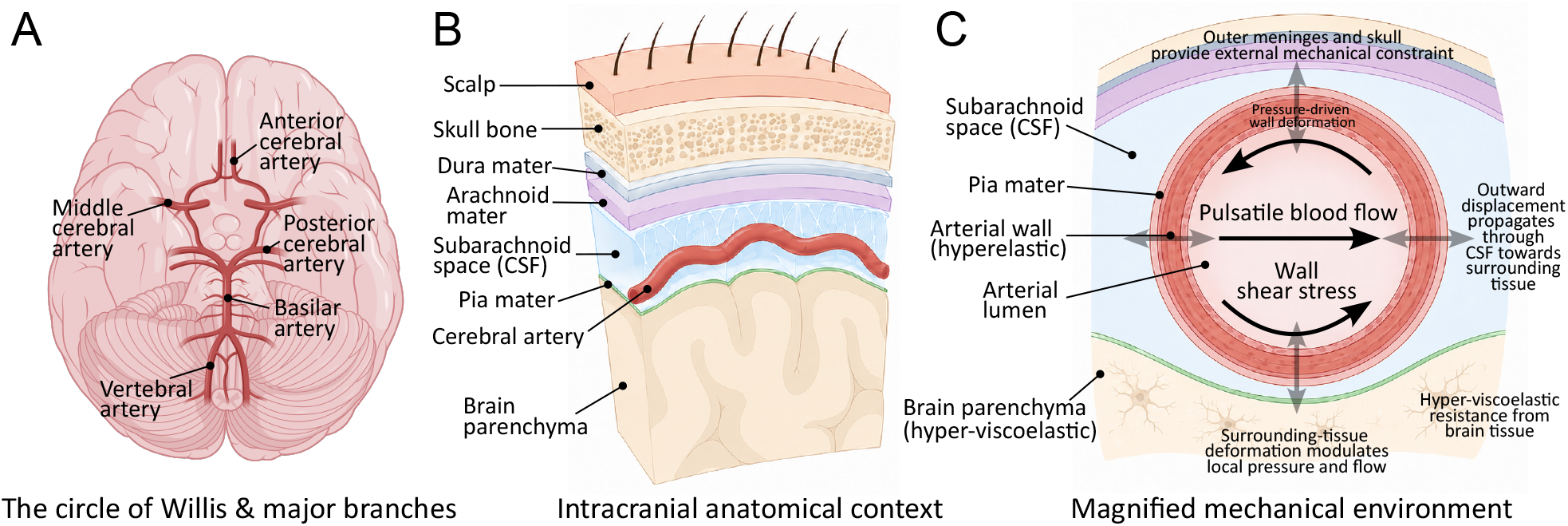
The circle of Willis and major cerebral arterial branches, generalised intracranial anatomical context, and mechanical coupling between a cerebral artery and the surrounding brain tissue. **A**: The circle of Willis and major cerebral arterial branches, including the anterior and posterior circulations, viewed from the inferior aspect of the brain. This subfigure was created in BioRender. **B**: Schematic oblique cutaway of the intracranial layers, displaying the scalp, skull, dura mater, arachnoid mater, CSF-filled subarachnoid space, pia mater, cerebral artery and underlying brain parenchyma. The artery is located within the subarachnoid space and follows the cortical surface. This schematic represents the general local environment of the modelled arterial segments rather than the exact anatomy of a specific vascular location. **C**: Magnified schematic of local blood–vessel–brain mechanical coupling. Pulsatile blood flow produces radial deformation of the compliant arterial wall. Vessel-wall motion is transmitted through the surrounding CSF towards the pia mater and adjacent hyper-viscoelastic brain parenchyma, while the tissue provides mechanical resistance to deformation. The outer meninges and skull impose additional external constraint. Deformation of the surrounding tissue can, in turn, modify the local mechanical environment of the vessel and flow.

Evidence that this coupling produces measurable tissue motion is provided by *in vivo* magnetic resonance imaging (MRI). Cardiac-resolved measurements have demonstrated sub-millimetre brain motion and measurable tissue strain throughout the cardiac cycle [11, 12, 13, 14, 15]. Displacement-encoding techniques, including DENSE MRI [16], have further shown that cardiac-induced deformation is spatially heterogeneous rather than a uniform expansion of the brain [17, 18]. Regional differences have been reported between cortical grey matter and deeper white matter, reflecting the local balance between vascular pulsatility and transient vascular volume change, tissue mechanical properties and surrounding mechanical constraint [11, 17, 18]. These observations establish vascular pulsatility as an internal mechanical driver of brain motion; however, imaging measurements alone cannot determine how much of the observed tissue response originates from local arterial geometry, vessel-wall deformation, tissue constitutive behaviour or the surrounding boundary conditions. Consequently, the mechanical pathway by which pulsatile haemodynamic loading is transmitted from an individual cerebral artery to the neighbouring brain tissue remains incompletely resolved.

This problem is further complicated by the heterogeneous mechanical behaviour of brain tissue. Magnetic resonance elastography (MRE) has demonstrated spatial variations in brain stiffness and systematic changes in apparent mechanical properties with ageing and neurological disease [19, 20, 21, 22, 23]. At the material level, brain tissue exhibits nonlinear and time-dependent behaviour arising from its cellular, extracellular and fluid constituents [22, 24, 25, 26, 27]. Under pulsatile loading, its response is therefore not purely elastic: mechanical energy is both stored and dissipated, producing frequency-dependent attenuation and phase differences between loading and deformation [28, 29, 30]. Regional variations in tissue properties may consequently alter not only the magnitude and spatial distribution of stress and strain, but also the transmission and dissipation of pulsatile mechanical energy through the blood–vessel–brain system, even under similar haemodynamic conditions [28, 31, 32, 33]. Although advanced MRI techniques can quantify subtle brain motion and strain non-invasively, their spatial resolution remains insufficient to resolve the local mechanics of individual blood–vessel–brain interactions or to isolate the respective contributions of vascular geometry, wall mechanics and tissue properties [13, 34]. More broadly, mechanistic studies of pressure-driven processes in the brain have demonstrated that fluid transport, tissue deformation and constitutive behaviour cannot always be treated independently, motivating coupled computational descriptions of brain–fluid interactions [35].

Computational modelling provides a complementary means of resolving these interactions by linking haemodynamics, vascular deformation and tissue mechanics within a common physical framework [1, 36, 33, 37, 38]. Nevertheless, most cerebral modelling studies have historically focused either on intravascular haemodynamics or on tissue-scale intracranial mechanics, while fully coupled representations of pulsatile blood flow, deformable cerebral arteries and surrounding brain tissue remain comparatively limited. In our previous work [10], we developed a coupled blood–vessel–brain finite-element framework incorporating non-Newtonian blood flow, hyperelastic arterial walls and hyper-viscoelastic brain tissue. That study demonstrated how blood pressure, heart rate, vessel dimensions and tissue properties can modulate pulse-driven brain deformation. However, the model employed an idealised axisymmetric geometry and was therefore designed primarily to identify fundamental mechanical relationships, without seeking to resolve the spatial effects introduced by realistic cerebral vascular anatomy.

This limitation becomes particularly important in anatomically complex cerebral arterial segments, where curvature, torsion and branching can generate strongly heterogeneous haemodynamic loading and vascular deformation. For example, the branching topology of the middle cerebral artery (MCA) and the pronounced curvature of the internal carotid artery (ICA) can produce marked spatial variations in pressure, wall shear and deformation that cannot be represented by an axisymmetric formulation [39, 40]. More generally, realistic arterial morphology may alter the local load transfer pathway by redistributing pulsatile forces along the vessel wall and into the surrounding tissue. Incorporating image-derived vascular anatomy is therefore necessary to determine how cerebral arterial geometry governs the spatial transmission of pulsatile mechanical loading across the blood–vessel–brain system.

In this study, we develop an image-derived three-dimensional (3D) computational framework to quantify pulsatile cerebral haemodynamics, arterial wall deformation and mechanically induced brain tissue motion within a unified finite-deformation fluid–structure interaction (FSI) formulation. Representative segments of the MCA, an MCA bifurcation, the basilar artery (BA) and the ICA are reconstructed from high-field 5 T MRI data acquired from a healthy adult subject. The framework couples non-Newtonian shear-thinning blood flow with hyperelastic cerebral arterial walls and hyper-viscoelastic brain tissue under physiologically representative pulsatile flow and pressure conditions. The predicted cardiac-induced tissue response is benchmarked against *in vivo* imaging measurements of brain deformation, providing an experimental reference for the magnitude and temporal characteristics of pulsatile tissue motion. The model is then used to determine how arterial geometry and wall thickness govern local vascular and tissue deformation, and how geometrical complexity influences near wall haemodynamic metrics. By integrating image-derived vascular anatomy, coupled blood–vessel–brain mechanics and *in vivo* benchmarking, this work establishes a mechanically consistent image-derived reference framework for cerebral pulsatility in the healthy brain and provides a foundation for quantifying how these interactions may be altered under pathological conditions.

The remainder of this paper is organised as follows. Section 2 describes the image-derived geometries, constitutive formulations, FSI implementation, physiological boundary conditions and post-processing procedures. Section 3 presents the *in vivo* benchmarking and compares haemodynamic and mechanical responses across the selected cerebral arterial segments. Section 4 discusses the biomechanical implications of geometry dependent blood–vessel–brain coupling and the relevance of the healthy reference framework to future pathological investigations. The main findings are summarised in Section 5.

## 2. Materials and methods

This section introduces the mathematical framework and the methodology adopted to investigate the coupled haemodynamic-biomechanical response of the cerebral vasculature and surrounding brain tissue. Firstly, the vasculature geometry reconstruction procedure is described. Subsequently, the mathematical framework governing blood flow, vessel wall mechanics, and brain tissue deformation is introduced, together with the fluid–structure interaction coupling strategy. The material properties and model parameters used in the simulations are then specified, followed by the model setup and applied boundary conditions. Then, the numerical implementation and computational procedures employed to solve the coupled system are described. Finally, the quantitative metrics used to evaluate the simulation outcomes are presented.

### 2.1. Vasculature reconstruction

The 3D geometrical models of four representative cerebral arteries, including the MCA (M1 segment), MCA bifurcation (M2 segment), ICA (C3-C7 segments) and basilar artery (BA), are reconstructed from the 5T time-of-flight magnetic resonance angiography (TOF-MRA) volume acquired from a healthy female participant in her 30s. Written informed consent is obtained from the participant, and the imaging protocol is approved by the institutional review board prior to MRI acquisition. Each image slice has a resolution of 602×736 pixels, with an in-plane pixel size of 0.3 mm and a slice thickness of 0.3 mm.

The arterial lumen is first segmented from the TOF-MRA dataset using intensity-based thresholding, with manual correction to preserve local vessel morphology, particularly in curved segments and bifurcations. Four representative arterial segments are then extracted, corresponding to anatomically distinct vascular configurations: a straight MCA segment, an MCA bifurcation, a curved BA segment, and a tortuous ICA segment, as shown in Figure 2. Following segmentation, triangulated surface meshes are generated and smoothed to remove stair-step artefacts arising from voxel discretisation while maintaining the physiological curvature and branch geometry of each vessel.

**Figure 2:**
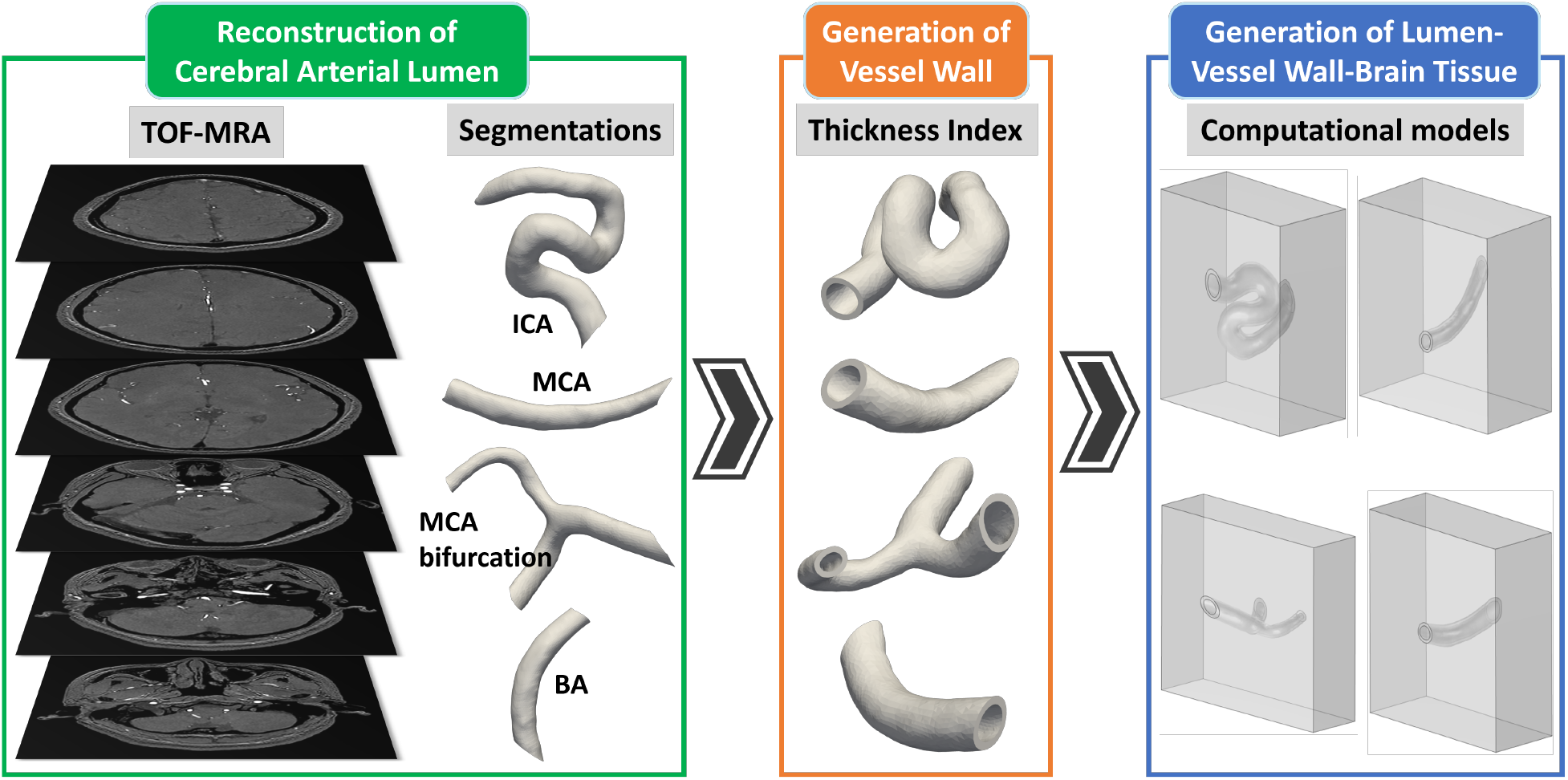
Schematic overview of the workflow used to reconstruct the cerebral arterial lumen geometries and generate computational lumen-vessel wall-tissue models. **Left panel (green)**: TOF-MRA images are used to reconstruct 3D geometries of cerebral arterial lumen through segmentation, including the ICA, MCA, MCA bifurcation and BA. **Middle panel (orange)**: Vessel wall geometries are generated from the segmented lumen by assigning thicknesses as listed in Table 1. **Right panel (blue)**: The resulting lumen–vessel wall–tissue models are incorporated into computational domains for subsequent simulations and analyses.

The segmented luminal surfaces serve as the inner arterial boundaries, and an outward offset operation is applied to generate the outer wall surface, enabling the biomechanical and FSI analyses of the reconstructed vessel wall models. A vessel wall thickness index is prescribed to ensure consistent wall generation along the vessel length while preserving local geometric continuity at bends and bifurcations. The mean, minimum and maximum vessel wall thickness measurements [41] across different arterial segments are summarised in Table 1. The resulting lumen and outer wall surfaces are then converted into vessel wall models for subsequent meshing and numerical simulation, which is displayed in Figure 2.

**Table 1:**
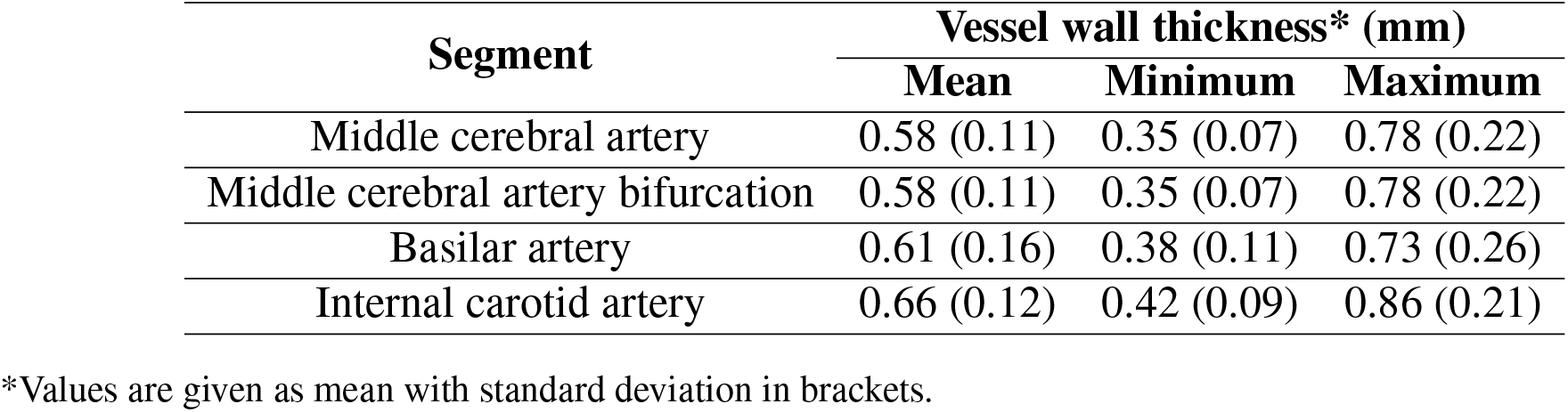
Vessel wall thickness of different arterial segments [41].

For each cerebral arterial segment, a surrounding solid domain is generated to represent the mechanical influence of the adjacent intracranial environment, as shown in Figure 2. This surrounding tissue region is defined by extending the outer vessel boundary with enough radial distance to avoid geometric interference while maintaining computational efficiency. Specifically, the extent of this surrounding domain is not intended to reproduce all anatomical layers adjacent to the artery, such as cerebrospinal fluid spaces, meninges, venous compartments, or fine perivascular interfaces. Instead, it serves as an effective continuum medium that enables consistent FSI analysis of arterial wall motion and deformation transfer into the adjacent environment.

On the other hand, the surrounding environment of the cerebral arterial segments is anatomically the Sylvian cistern rather than direct bulk parenchyma. Available anatomical studies [42] indicate that the immediate Sylvian subarachnoid compartment is on the order of 1 mm, whereas the adjacent insular/opercular parenchymal boundaries lie at substantially greater depths, commonly around 18–25 mm and extending up to 35 mm in the posterior insular region. Accordingly, the outer domain dimensions were selected to provide sufficient separation between the vessel wall and the external boundaries, thereby minimising the influence of the imposed boundary constraints on local deformation patterns, while maintaining geometric consistency across all four models without incurring unnecessary computational cost. Finally, Boolean operations are then performed to ensure conformal interfaces between the vessel wall and surrounding tissue domains, enabling coupled structural analysis of vessel–tissue interaction.

### 2.2. Mathematical model

#### 2.2.1. Governing equations of blood flow

Blood flow in the lumen is modelled as laminar flow of an incompressible, shear-thinning non-Newtonian fluid [43], with a Reynolds number in the reported range of 300 to 700 [44]. The governing equations are the conservation of linear momentum and mass, written as:

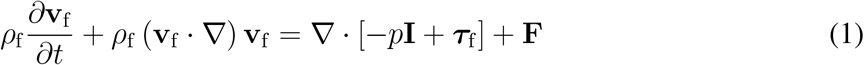

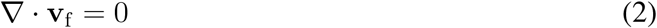

where *ρ*_f_ is the constant blood density, **v**_f_ is the blood velocity vector, *p* is the pressure, **I** is the identity tensor, ***τ***_f_ is the viscous stress tensor and **F** denotes the body force term, such as gravity, which is taken to be 0 in the present study. For the present model, the viscous stress tensor is defined as:

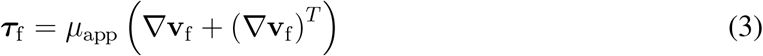

where *µ*_app_ is the apparent viscosity of blood. Unlike a Newtonian fluid, the viscosity here is not constant, but varies with the local shear rate. The apparent viscosity is described using the Carreau shear-thinning constitutive model [45]:

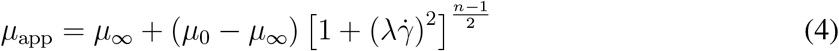

where *µ*_0_ and *µ*_∞_ are the zero-shear and infinite-shear viscosities, respectively, *λ* is the characteristic time constant, n is the flow behaviour index and 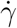 is the magnitude of the shear rate, defined as:

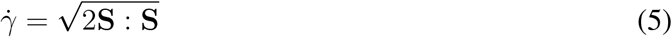

where **S** is the strain-rate tensor:

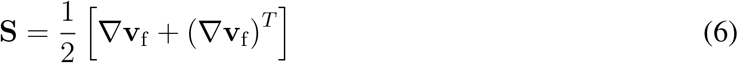

and

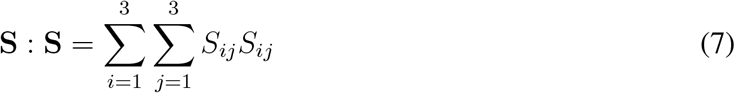

where *S*_*ij*_ is the component of the strain-rate tensor. Therefore, 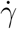 represents the scalar magnitude of the local rate of deformation.

In the Carreau model, *µ*_0_ *> µ*_∞_ *>* 0, *λ >* 0 and 0 *< n <* 1. Therefore, the apparent viscosity *µ*_app_ decreases from *µ*_0_ towards *µ*_∞_ as the shear rate 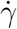 increases, thereby representing the shear-thinning behaviour of blood.

#### 2.2.2. The interaction between blood flow and vessel wall

The coupling between blood flow and vessel wall deformation is described using a two-way FSI scheme. At the blood–vessel interface, continuity of velocity, displacement and traction equilibrium is imposed to ensure the kinematic and dynamic compatibility between the fluid and solid domains. The continuity of displacement and velocity at the interface can be written as:

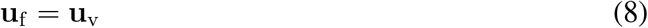

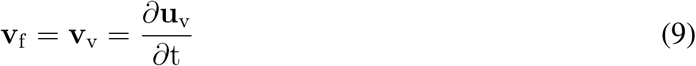

where **u**_f_ denotes the displacement of the fluid mesh at the interface and **u**_v_ is the displacement of the vesell wall. The traction equilibrium condition is given by:

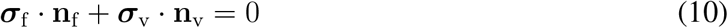

where ***σ***_f_ and ***σ***_v_ are the Cauchy stress tensors in the fluid and vessel-wall domains, respectively. **n**_f_ and **n**_v_ are the outward unit normal vectors of the fluid and solid domains, respectively. Because the vessel deforms under pulsatile blood loading, the boundary of the fluid domain evolves with time. To account for the moving boundary induced by vessel wall deformation, the Arbitrary Lagrangian–Eulerian (ALE) framework [46] is adopted. At the blood–vessel interface, the mesh velocity is given by combining the displacement continuity and no-slip conditions, as:

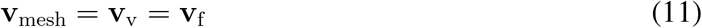

Under this description, the convective term in the fluid momentum equation is evaluated relative to the mesh velocity, and Eq. (1) can therefore be rewritten in the ALE frame as:

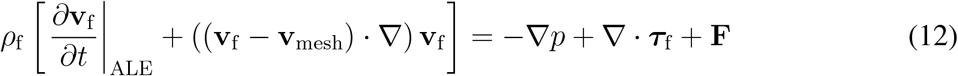

The ALE formulation enables consistent transmission of pressure and shear loading from the blood to the deformable vessel wall while preserving mesh conformity at the fluid-solid interface throughout the pulsatile loading cycle.

#### 2.2.3. Governing equations of vessel wall movement

The vessel wall is modelled as a deformable hyperelastic solid undergoing finite deformation [47, 48, 49, 50]. Its motion is governed by the balance of linear momentum in the reference configuration, as:

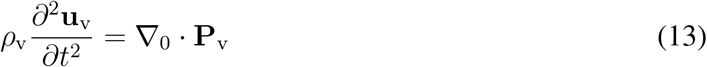

where *ρ*_v_ is the density of the vessel wall, ∇_0_ denotes the gradient operator with respect to the reference configuration, and **P**_v_ is the first Piola-Kirchhoff stress tensor.

The deformation gradient is defined as:

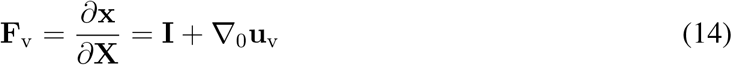

where **X** and **x** are the coordinates in the reference and current configurations, respectively.

The constitutive response of the vessel wall is obtained from a strain-energy density function *W*_v_, such that the first Piola–Kirchhoff stress is obtained by differentiating *W*_v_ with respect to the deformation gradient [47, 48, 49, 50]:

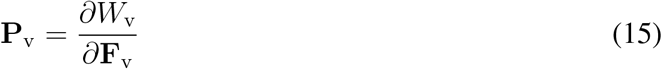

The corresponding Cauchy stress tensor is obtained through the standard Piola transformation [47, 48], as:

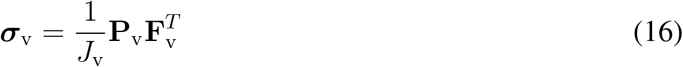

where *J*_v_ = det(**F**_v_) is the local volume ratio.

To capture the nonlinear elastic response of the arterial wall, a one-term Ogden hyperelastic model is adopted [37]. The strain-energy density function is expressed as:

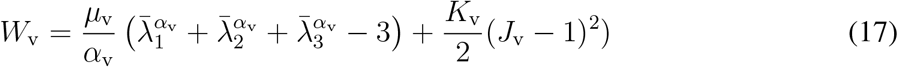

where *µ*_v_ and *α*_v_ are the material parameters that represent the shear modulus and the type of nonlinearity, respectively. *K*_v_ is the bulk modulus. 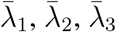 are the principal isochoric stretches.

#### 2.2.4. Governing equations of brain tissue deformation

Following our previous study, where we implemented a hyper-viscoelastic model in the COMSOL Multiphysics platform by incorporating Storakers hyperelasticity and generalised Maxwell viscoelasticity [10], the surrounding brain tissue is modelled as an isotropic finite-strain hyper-viscoelastic material to account for its nonlinear instantaneous deformation and subsequent time-dependent stress relaxation [51, 52, 53, 54]. The motion of the brain tissue satisfies the balance of linear momentum in the solid domain:

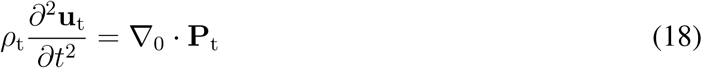

where *ρ*_t_ is the tissue density, **u**_t_ is the tissue displacement vector, and **P**_t_ is the first Piola-Kirchhoff stress tensor in the brain domain.

The finite-deformation kinematics are described by the deformation gradient, as:

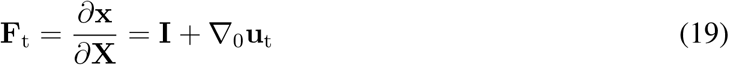

with the corresponding volume ratio:

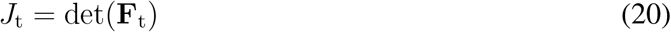

While Eq. (19) is adopted to describe the finite deformation of brain tissues, their total deformation gradient **F**_t_ is multiplicatively decomposed into elastic and viscous components:

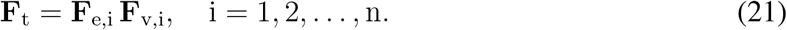

where **F**_e,i_ and **F**_v,i_ are the elastic and viscous components associated with the i-th Maxwell branch.

The Helmholtz free-energy density of brain tissue is then written as the sum of equilibrium and non-equilibrium contributions:

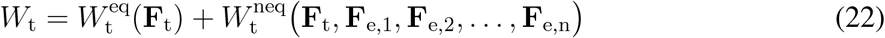

where 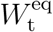 and 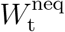 represents the time-independent hyperelastic equilibrium response and time- and rate-dependent viscoelastic non-equilibrium response, respectively.

The non-equilibrium contribution is further decomposed as:

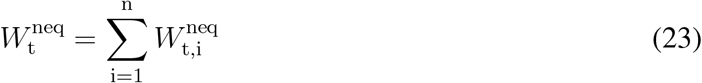

where 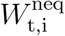 is the free-energy contribution of the i-th viscoelastic branch. This branch-wise decomposition follows the generalised Maxwell rheological framework. For the equilibrium response, a one-term Storakers hyperelastic model is employed and written as [48, 55]:

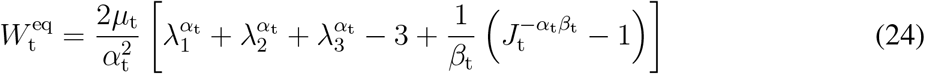

where *µ*_t_ is the shear modulus, *α*_t_ is the nonlinearity parameter, *λ*_1_, *λ*_2_ and *λ*_3_ are the principal stretches, and *β*_t_ controls the volumetric response.

Following Eq. (15) and substituting Eq. (22), the total first Piola-Kirchhoff stress is correspondingly decomposed into equilibrium and non-equilibrium parts:

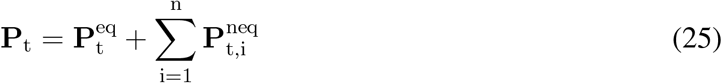

where

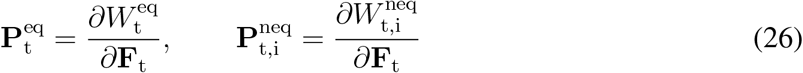

To represent viscous relaxation, the overstress in each Maxwell branch evolves with its own characteristic relaxation time. In compact form, the branch response may be represented through a Prony-series-type description:

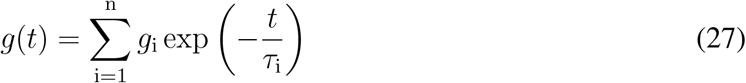

where *g*_i_ is the dimensionless energy factor, also interpreted as the relative modulus of the i-th Maxwell branch, and *τ*_i_ is the corresponding relaxation time. Accordingly, the time-dependent shear modulus can be expressed as:

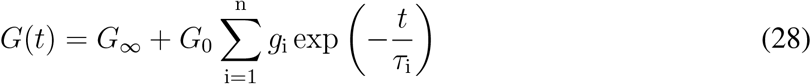

where the instantaneous shear modulus is denoted by *G*_0_ = *µ*_t_ and the long-term shear modulus is calculated as:

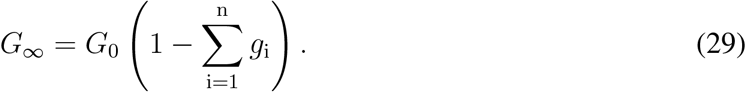

The non-equilibrium stresses associated with the Maxwell branches are evaluated from the specific Prony-series parameters *g*_i_ and *τ*_i_. The resulting constitutive structure allows the model to capture the rate-dependent compression-relaxation behaviour of brain tissues.

The corresponding Cauchy stress tensor in the tissue domain is obtained as:

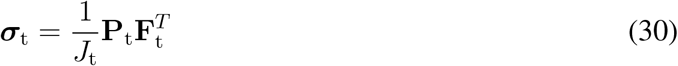

Therefore, the total brain-tissue response is governed by the combination of the instantaneous hyperelastic contribution, the long-term equilibrium response, and the time-dependent overstresses carried by the viscoelastic Maxwell branches.

### 2.3. Model parameters

The material and constitutive parameters used in the FSI model are summarised in Table 2, and their sources are provided accordingly. These include the density and Carreau-model parameters of blood, the hyperelastic parameters of the vessel wall, and the hyper-viscoelastic parameters of the surrounding brain tissue. The hyper-viscoelastic material properties of the human brain tissues are derived from our previous experimental results. Readers can refer to our previous study for further experimental details and the derivations of brain tissue properties [10, 32].

**Table 2:** Material and constitutive parameters used in the mathematical model.

| Parameter | Symbol | Unit | Value | Source |
| --- | --- | --- | --- | --- |
| Blood density | $\rho_f$ | kg/m<sup>3</sup> | 1050 | [56] |
| Zero-shear viscosity | $\mu_0$ | Pa · s | 0.056 | [56, 57] |
| Infinite-shear viscosity | $\mu_\infty$ | Pa · s | 0.0035 | [56, 57] |
| Carreau relaxation time | $\lambda$ | s | 3.313 | [56, 57] |
| Power-law index | $n$ | – | 0.3568 | [56, 57] |
| Vessel wall density | $\rho_v$ | kg/m<sup>3</sup> | 1000 | [37, 58] |
| Ogden shear modulus | $\mu_v$ | Pa | $2.0 \times 10^5$ | [37, 59] |
| Ogden exponent | $\alpha_v$ | – | 2 | [37, 59] |
| Bulk modulus of vessel wall | $K_v$ | Pa | $1.0 \times 10^8$ | [37, 59] |
| Brain tissue density | $\rho_t$ | kg/m<sup>3</sup> | 1050 | [60, 61] |
| Instantaneous shear modulus | $G_0$ | Pa | 624.11 | [10, 32] |
| Long-term shear modulus | $G_\infty$ | Pa | 64.43 | [10, 32] |
| Storakers exponent | $\alpha_t$ | – | -3.3273 | [10, 32] |
| Storakers compressibility parameter | $\beta_t$ | – | 49 | [10, 32] |
| Energy factor | $g_i$ | – | (0.600, 0.048, 0.249) | [10, 32] |
| Relaxation time | $\tau_i$ | s | (1.218, 35.922, 971.53) | [10, 32] |

### 2.4. Model setup and boundary conditions

As shown in Figure 3A, the vessel lumen, vessel wall and surrounding tissue are explicitly represented for each computational domain. It should be noted that the surrounding tissue domain is partitioned into three concentric layers adjacent to the vessel and a remaining exterior region. The layer boundaries are generated using an in-house Python algorithm and conformed to the geometry of the embedded cerebral artery, with each successive layer extending further into the surrounding tissue. Quantitative post-processing is restricted to these three layers to reduce the influence of boundary-condition artefacts arising from constraints applied to the outer surfaces of the cuboidal tissue domain. The resulting strain fields therefore provide a more representative measure of the local tissue deformation induced by vascular pulsation. The volume of the third tissue layer is selected such that the ratio of vascular volume to local tissue volume is comparable to the reported cerebral blood volume fraction of approximately 4–6% in grey matter [62]. This criterion is used to define a physiologically representative tissue volume surrounding each arterial segment.

**Figure 3:**
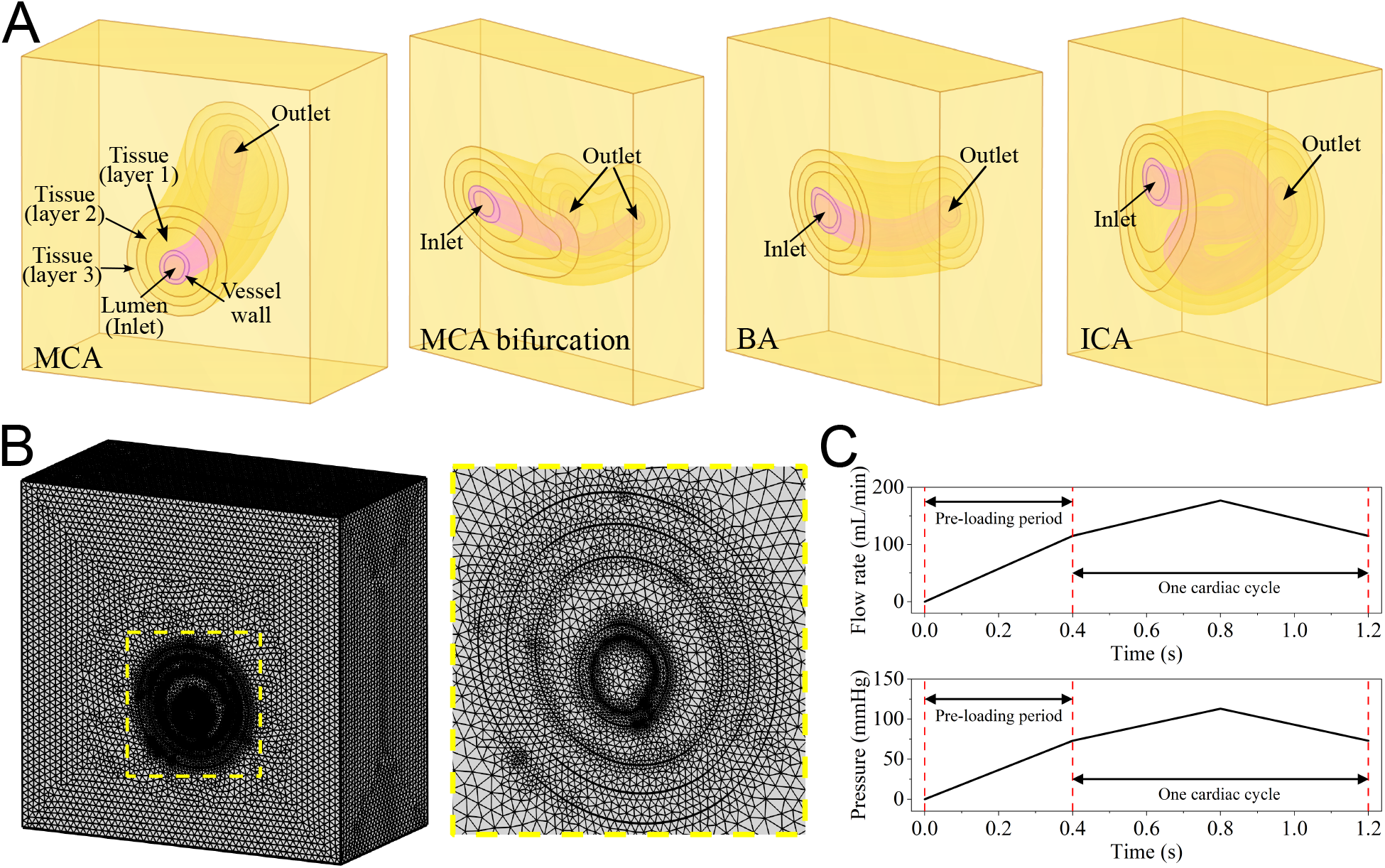
Computational model setup and prescribed boundary conditions. **(A)** Each vessel is embedded within a surrounding tissue domain, with the vessel lumen, vessel wall, and surrounding brain tissue explicitly represented. For quantitative post-processing, volumetric strain is extracted only from the three surrounding tissue layers, rather than from the entire cuboid tissue block, to avoid artificial strain effects caused by constrained outer tissue boundaries. Inlet and outlet boundaries are indicated for each computational model. **(B)** Representative computational mesh of MCA and zoom-in view near the vessel and surrounding tissue layers. **(C)** Time-dependent inlet flow rate and outlet pressure profiles are used to apply physiological pulsatile loading on MCA. The interval before 0.4 s represents a pre-loading ramp-up period, during which the applied flow rate and pressure are gradually increased to improve numerical stability. After this period, the physiological pulsatile loading in a cardiac cycle is applied. Segment-specific flow rates and systolic/diastolic pressures are listed in Table 3.

All geometrical domains are imported into the COMSOL Multiphysics 6.4 platform for the inspection of surface continuity, topological consistency and grid convergence prior to computational analysis. Figure 3B shows the representative finite element mesh of the MCA model and its zoomed-in view near the vessel and surrounding tissue layers. The finest mesh with an averaged element size of 0.56 mm is applied in the area near the interface between the vessel wall and the surrounding brain tissue to achieve high-fidelity modelling results.

**Table 3:** Segment-specific blood flow rate and blood pressure.

| Segment | Blood flow rate<br>(mL/min) [64] | Blood pressure (mmHg) [63] |  |
| --- | --- | --- | --- |
|  |  | Systolic pressure | Diastolic pressure |
| Middle cerebral artery | $146 \pm 31$ | 113 | 73 |
| Middle cerebral artery bifurcation | $146 \pm 31$ | 113 | 73 |
| Basilar artery | $145 \pm 41$ | 113 | 73 |
| Internal carotid artery | $257 \pm 48$ | 117 | 77 |

Physiologically relevant boundary conditions are prescribed for the blood, vessel wall, and surrounding brain tissue domains. For each arterial segment, a pulsatile inlet velocity is imposed based on the corresponding volumetric flow rate, and a pulsatile outlet pressure is defined using the systolic and diastolic pressures of each segment [63], as listed in Table 3. The inlet velocity is calculated from the imposed flow rate [64] and the inlet cross-sectional area. To avoid abrupt loading and improve numerical stability, both the inlet velocity and outlet pressure are ramped from zero to their prescribed values over an initial pre-loading period of 0.4 s. After this, the physiological pulsatile loading in a cardiac cycle is applied. The time-dependent inlet flow rate and outlet pressure profiles are plotted in Figure 3C. No-slip boundary condition is imposed at the blood-vessel interface.

The blood-vessel wall interface is set as a two-way FSI interface, such that continuity of displacement and traction is satisfied between the fluid and vessel wall domains, as described in Section 2.2.2. The outer vessel wall is bonded to the surrounding brain tissue, and continuity of displacement and traction is also imposed at the vessel-tissue interface.

For the brain tissue domain, the outer boundary of the cuboid tissue block is fixed to represent the mechanical support provided by the surrounding intracranial environment, as described in Figure 1B. Because this constraint may artificially influence the deformation field near the outer boundary, as stated before, quantitative analysis is restricted to the three tissue layers surrounding the vessel, which is described in detail in Section 3.1.

### 2.5. Numerical implementation

The coupled blood-vessel-brain model is implemented in COMSOL Multiphysics 6.4 using the finite element method (FEM). The blood domain is formulated in the ALE framework [46], while the vessel wall and brain tissue are described in a Lagrangian material framework. Deformation of the fluid domain is accommodated using a moving mesh, whose displacement is coupled to that of the fluid–solid interface to maintain mesh conformity as the vessel deforms over the cardiac cycle. A fully coupled solver is employed to solve the strongly coupled system of blood flow, vessel wall deformation, and brain tissue dynamics.

### 2.6. Quantification of outcomes

The simulation outcomes are quantified in terms of haemodynamic characteristics and cardiac-induced brain tissue deformation. The quantitative indices are described below.

#### (1) Volumetric strain of brain tissues

The volumetric strain quantifies the local relative volume change of the vessel wall and surrounding brain tissue. Since both solid domains are formulated within a finite-deformation framework, the volumetric strain is evaluated from the determinant of the deformation gradient as [47]:

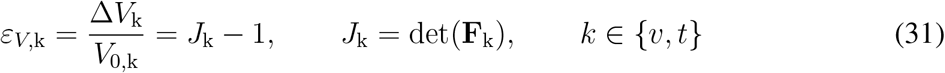

where Δ*V*_k_ = *V*_k_ − *V*_0,k_, *V*_0,k_ and *V*_k_ denote the volumes of a material element in the reference and current configurations, respectively. The subscripts v and t refer to the vessel wall and brain tissue. A positive value of *ε*_*V*,k_ indicates local volumetric expansion, whereas a negative value indicates local compression.

#### (2) Time-averaged wall shear stress

The time-averaged wall shear stress (TAWSS) quantifies the average magnitude of the shear traction exerted by the blood flow on the luminal surface of the cerebral arteries over one cardiac cycle, defined as [65, 66]:

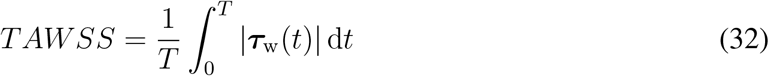

where *T* is the duration of one cardiac cycle, ***τ***_w_(*t*) is the instantaneous WSS vector.

#### (3) Oscillatory shear index

The oscillatory shear index (OSI) quantifies the temporal variation in the direction of the WSS vector over one cardiac cycle, defined as [65, 66]:

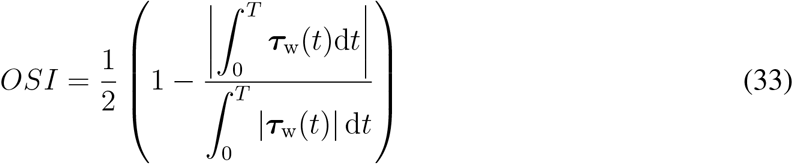

The OSI ranges from 0 to 0.5. A value close to 0 indicates predominantly unidirectional WSS, whereas a value approaching 0.5 indicates strong directional oscillation or reversal during the cardiac cycle.

#### (4) Relative residence time

The relative residence time (RRT) is a derived haemodynamic index that combines the effects of low TAWSS and directional oscillation of the WSS vector, which is defined as [65, 66]:

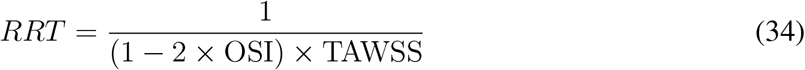

Higher RRT values indicate regions characterised by low-magnitude and/or highly oscillatory WSS and may therefore be used as a surrogate marker of prolonged near-wall particle residence or flow stagnation.

## 3. Results

This section presents the simulation results. First, experimentally measured pulsatile brain deformation available in the literature is used to validate the predictive accuracy of the framework. Then, the pulsatile blood flow characteristics and the associated deformation of the vessel wall and brain tissue are examined under physiological flow conditions. The influence of vessel wall thickness on the coupled blood–vessel–brain dynamics is also investigated to evaluate how structural variations in the arterial wall affect flow behaviour, wall motion and tissue deformation. Finally, the haemodynamic responses in the vicinity of the vessel wall are analysed through three key vessel wall-based indices, i.e., TAWSS, OSI and RRT, to further assess the spatial characteristics of near-wall flow in different cerebral artery models.

### 3.1. Model validation

The proposed computational framework is first validated by comparing the simulated volumetric strain in the surrounding brain tissue with experimentally measured volumetric strain over a cardiac cycle reported in Ref [17, 18]. In this section, all geometrical models are constructed using the mean vessel wall thickness for validation, as listed in Table 1. In our previous 2D study [10], simulations over two cardiac cycles produced essentially identical results for both cycles; therefore, in the present study, only one cardiac cycle is simulated and used for the subsequent analysis. The temporal evolution of simulated volumetric strain over one cardiac cycle (0.4 to 1.2 s) is then employed and compared against the reported experimental range for brain grey matter [17], as shown in Figure 4A. The simulated results are normalised relative to the value at the simulation time of 0.4 s, such that the pre-loading deformation is removed and subsequent strain values represent changes occurring after the pre-loading period. Among the three computational models, the volumetric strain increases during systole, reaches a peak near the systolic phase of the cardiac cycle, at approximately 30% of the cycle, and subsequently decreases during diastole. This temporal pattern is consistent with the experimentally observed cardiac-cycle-dependent variation in brain tissue deformation.

**Figure 4:**
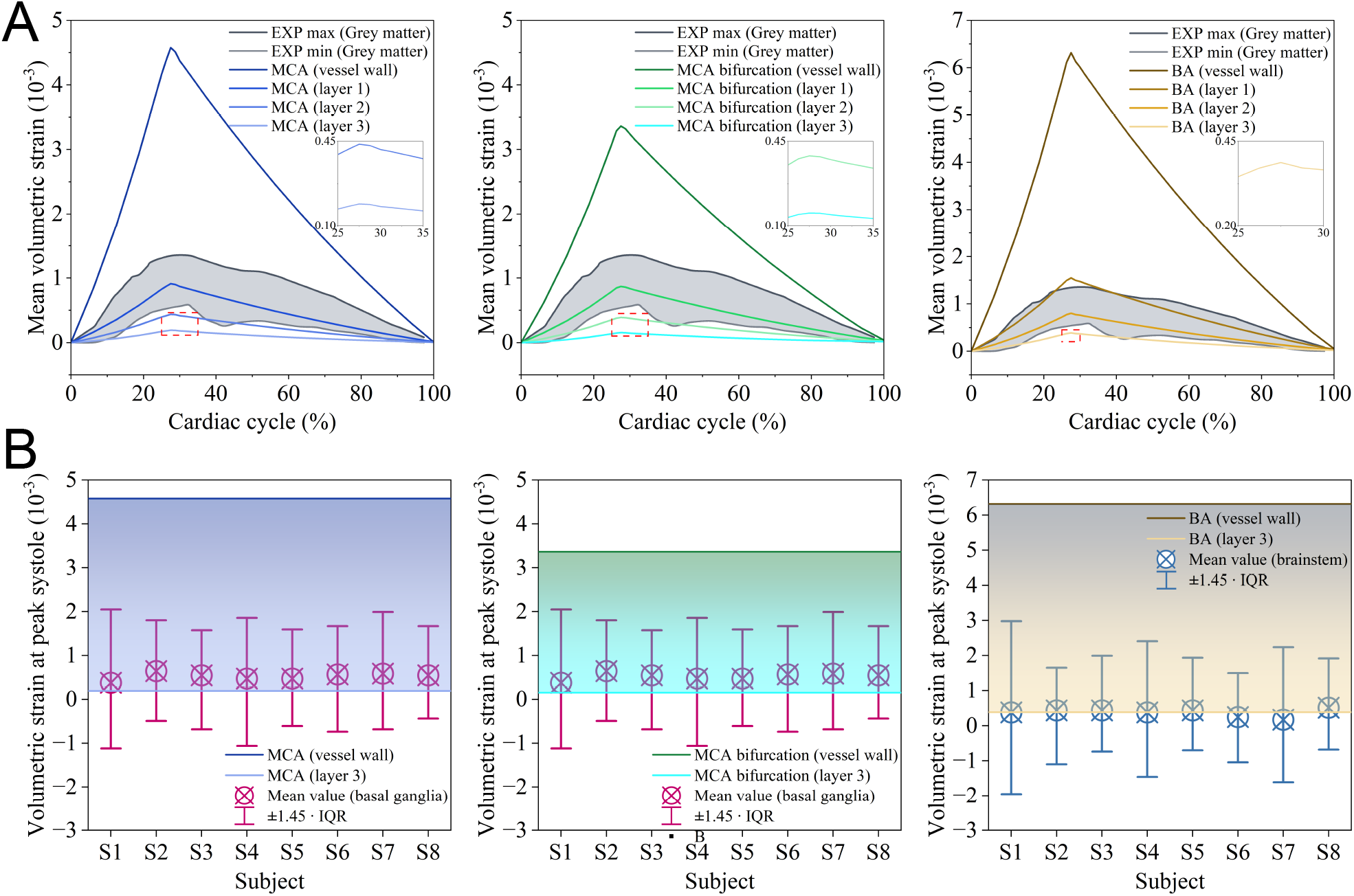
Validation of the blood–vessel–brain dynamics model against reported experimental results obtained by MRE. **(A)** Comparison of the simulated temporal evolution of volumetric strain over one cardiac cycle with the experimentally measured range reported in [17]. For each model, volumetric strain of the vessel wall and the three nested perivascular tissue layers is estimated, with each successive region incorporating the preceding inner regions. The grey-shaded area represents the experimentally measured range bounded by the minimum and maximum values of the brain grey matter. The inset subplots provide enlarged views of the low-amplitude strain response in the third tissue layer near peak systole. **(B)** Subject-wise comparison of volumetric strain at peak systole of the cardiac cycle. The shaded area shows the simulated volumetric strain at peak systole. Its upper boundary represents the result of the vessel wall, and its lower boundary represents the results of the tissue within layer 3. The bars show experimentally measured subject-specific mean strains, with error bars representing the non-parametric reproducibility coefficient (*±*1.45*×*IQR) [18]. Basal-ganglia measurements are used for the MCA and MCA-bifurcation models, while brainstem measurements are used for the BA model.

In the present study, volumetric strain over a cardiac cycle is extracted from three surrounding tissue layers. The rationale for selecting these tissue layers is described in Section 2.4. The regions are defined cumulatively, such that each successive region includes the vessel wall and all preceding inner tissue regions, thereby representing a progressively larger perivascular volume. As expected, the simulated strain is largest in the vessel wall, and gradually decreases toward the outer tissue layers, as shown in Figure 4A. This reduction reflects both the attenuation of vessel-induced deformation with distance from the vessel and the averaging of the response over an increasingly larger tissue volume. Therefore, strain values of the vessel wall are not expected to fall within the experimentally reported grey matter range. This is because the vessel wall represents a highly localised near-lumen region, where deformation is directly driven by the blood flow and exceeds voxel- or region-averaged experimental measurements. In contrast, the progressively enlarged tissue regions incorporate increasing amounts of tissue farther from the vessel, where the transmitted deformation is attenuated.

The experimentally measured strain of brain grey matter is therefore used as a physiological reference for the overall magnitude and temporal behaviour of tissue deformation, rather than as a strict layer-by-layer validation criterion. For the MCA, MCA bifurcation, and BA models, the simulated tissue-layer responses generally overlapped with, or bounded, the experimentally reported range during most of the cardiac cycle, as shown in Figure 4A. Among the simulated cases, the BA model exhibits the largest peak tissue strain, followed by the MCA and its bifurcation model, reflecting the combined effects of vessel geometry, local vascular configuration, blood flow rate, and pressure magnitude on the deformation transmitted to the surrounding tissue. The ICA model is excluded from the quantitative validation in this section to avoid anatomically inconsistent comparisons, as no directly corresponding experimental volumetric strain measurements are available for ICA-adjacent tissue. However, the ICA model is still included in the subsequent computational analysis.

To further validate the model, the simulated volumetric strain at peak systole is compared against subject-specific measurements in another experimental study [18], as shown in Figure 4B. According to the relevant anatomical regions, simulated values from the MCA and MCA bifurcation models are compared with basal ganglia measurements, while those from the BA model are compared with brainstem measurements. The experimental data are presented by subject-specific mean values from two repeated scans among 8 subjects, together with the non-parametric reproducibility coefficient, represented by the error bar and defined as ±1.45 × interquartile range (IQR) [17]. The simulated volumetric strain range between the vessel wall and the third tissue layer covers the transition from the highly localised vessel-adjacent response to the attenuated deformation in the outer tissue layer. Most simulated volumetric strain ranges fall within or near the experimental measurements. Overall, these comparisons suggest that the proposed computational framework can reproduce both the magnitude and cardiac-cycle-dependent variation of experimentally reported brain volumetric strain, thereby demonstrating the predictive accuracy of the developed model in this study.

### 3.2. The pulsatile blood flow and tissue deformation

Following model validation, the spatiotemporal distribution of volumetric strain is evaluated over one cardiac cycle to investigate how arterial pulsation-induced deformation is transmitted from the vessel wall to the surrounding brain tissue. Figure 5A-D shows the volumetric strain distributions at selected time points from 0.4 to 1.2 s for the MCA, MCA bifurcation, BA, and ICA models, respectively. The time point of 0.4 s corresponds to the end of the initial pre-loading period, after which the physiological pulsatile flow and pressure loading are applied, as depicted in Figure 3C. Positive volumetric strain represents local tissue expansion, whereas negative volumetric strain represents local compression. Mean vessel wall thickness is used in this section.

**Figure 5:**
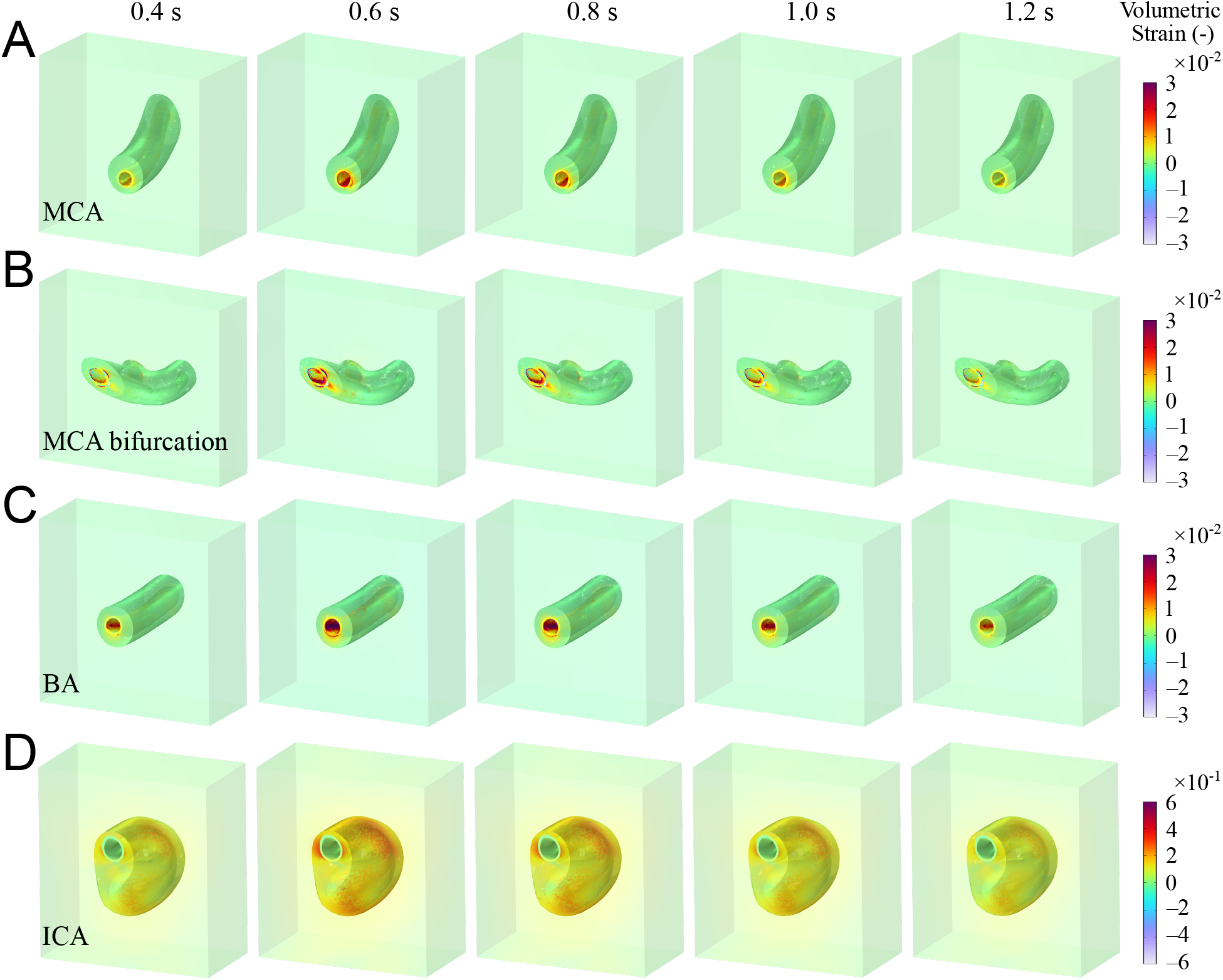
Temporal evolution of volumetric strain over one cardiac cycle. Volumetric strain distributions are shown at five typical time points after the initial pre-loading period, from 0.4 to 1.2 s, for the **(A)** MCA, **(B)** MCA bifurcation, **(C)** BA and **(D)** ICA models. All models shown use the mean-thickness configuration. For each model, volumetric strain is visualised in the vessel wall and the surrounding tissue domain to illustrate the transmission of cardiac-pulsation-induced deformation from the vessel to the adjacent brain tissue. Only the first tissue layer is extracted from the cuboid tissue block for visualisation here. Positive volumetric strain indicates local expansion, whereas negative volumetric strain indicates local compression. Note that different colour scales are used for the ICA model due to its larger volumetric strain magnitude. For better visualisation, the first tissue layer is rendered with a transparency of 0.3, while the remaining surrounding tissue is rendered with a transparency of 0.5.

Across all models, the volumetric strain is concentrated near the vessel wall and gradually attenuates with distance into the surrounding tissue domain, as displayed in Figure 5. This indicates that cardiac pulsation-induced deformation is highly localised around the cerebral arteries, with the largest deformation occurring within the vessel wall and immediately adjacent tissue. Following the pre-loading period, volumetric strain increases with the hydraulic load, reaches its maximum during systole, and subsequently decreases during diastole. This temporal behaviour is consistent with the prescribed pulsatile inlet flow rate and outlet pressure profiles.

The MCA model shows a localised strain concentration around the vessel lumen, with deformation transmitted primarily to the surrounding tissue near the curved arterial segment, as displayed in Figure 5A. The MCA bifurcation model (Figure 5B) exhibits a more spatially heterogeneous strain distribution because of its more complex vascular geometry. In particular, strain concentration is observed around the bifurcation and near the inlet region, suggesting that local curvature and branching can influence the distribution of deformation transmitted to the adjacent tissue. The BA model shows a relatively symmetric strain pattern around the vessel wall, which is plotted in Figure 5C. This is consistent with its simpler tubular geometry. Compared with the MCA bifurcation, the deformation in the BA model is more localised around the vessel and shows less spatial heterogeneity. The ICA model exhibits a larger and more spatially extended volumetric strain response than the other models. Results indicate that the ICA can transmit stronger pulsation-induced deformation to the surrounding tissue domain. This difference is reflected by the different colour scale used for the ICA results, as shown in Figure 5D. The larger magnitude of volumetric strain can be attributed to the tortuous geometry and higher inlet flow and outlet pressure loading applied to this model. In addition, the ICA serves as a major upstream feeding source that supplies blood to the anterior and middle cerebral territories. As a result, the total blood flow carried by the ICA is substantially greater than that in an individual downstream MCA branch or in the posterior circulation represented by the BA, as listed in Table 3.

Figure 6 complements the volumetric strain results by showing the displacement magnitude of the vessel wall and adjacent tissue. In general, the displacement follows the temporal variation of the cardiac loading, i.e., increasing during systole and decreasing during diastole. Unlike volumetric strain, which describes local expansion and compression, displacement represents the overall motion transmitted from the pulsating vessel to the surrounding tissue.

**Figure 6:**
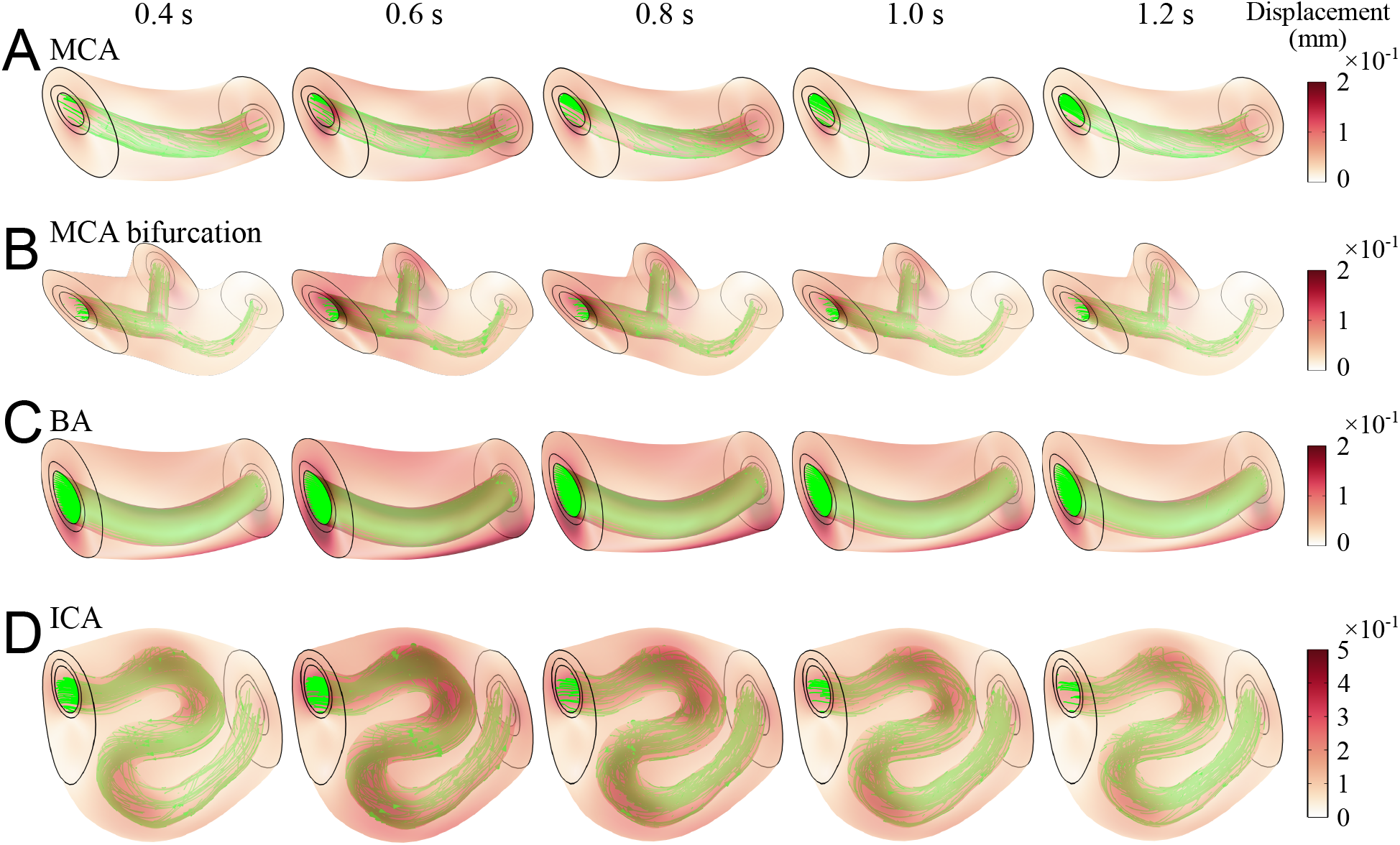
Temporal evolution of tissue displacement and flow condition over one cardiac cycle. Displacement distributions are shown at five typical time points after the initial pre-loading period, from 0.4 to 1.2 s, for the **(A)** MCA, **(B)** MCA bifurcation, **(C)** BA and **(D)** ICA models. All models shown use the mean-thickness configuration. For each model, displacement magnitude is visualised in the vessel wall and the first tissue layer to illustrate the transmission of cardiac-pulsation-induced deformation from the vessel to the adjacent brain tissue. The green lines within the vessel lumen represent flow velocity streamlines, while the displacement maginitude is indicated by the colour contours. A different colour scale is used for the ICA model because of its larger displacement magnitude. For better visualisation, the first tissue layer is rendered with a transparency of 0.5.

The MCA model shows a relatively localised and smooth displacement distribution around the curved vessel segment (Figure 6A), whereas the MCA bifurcation exhibits a more heterogeneous pattern around the inlet and branching region (Figure 6B). The BA model displays a comparatively uniform displacement field, consistent with its simpler tubular geometry (Figure 6C). In contrast, the ICA produces the largest and most spatially extensive displacement response, as reflected by the higher colour scale used in Figure 6D. This stronger response is associated with its tortuous geometry and greater haemodynamic loading. Overall, the displacement results are consistent with the volumetric strain patterns in Figure 4, demonstrating that vessel geometry and loading conditions strongly influence the magnitude and spatial extent of pulsation-induced deformation transmitted to the surrounding brain tissue.

### 3.3. Effect of vessel wall thickness on the blood-vessel-brain dynamics

As shown in Table 1, vessel wall thickness varies within a defined range. While the mean wall thickness of each arterial segment is used in Sections 3.1 and 3.2, this section examines the sensitivity of the mean volumetric strain over the cardiac cycle to variations in wall thickness. The analysis is performed for the MCA, MCA bifurcation, BA, and ICA models, with the results presented in Figure 7A–D. In all models, the volumetric strain increases during systole, reaches its maximum at approximately 30% of the cardiac cycle, and subsequently decreases toward zero by the end of the cycle. A consistent inverse relationship over all models is observed between wall thickness and volumetric strain: the minimum-thickness models produce the highest volumetric strain, whereas the maximum-thickness models exhibit the lowest volumetric strain. Under the same pulsatile loading condition, a thinner vessel wall has lower structural stiffness and carries the applied load over a smaller cross-sectional area, resulting in higher wall stresses and greater deformation transmitted to the surrounding tissue. Conversely, increasing the thickness of the vessel wall distributes the applied load over a larger wall cross-section and increases the overall structural stiffness of the vessel, thereby reducing the volumetric strain.

**Figure 7:**
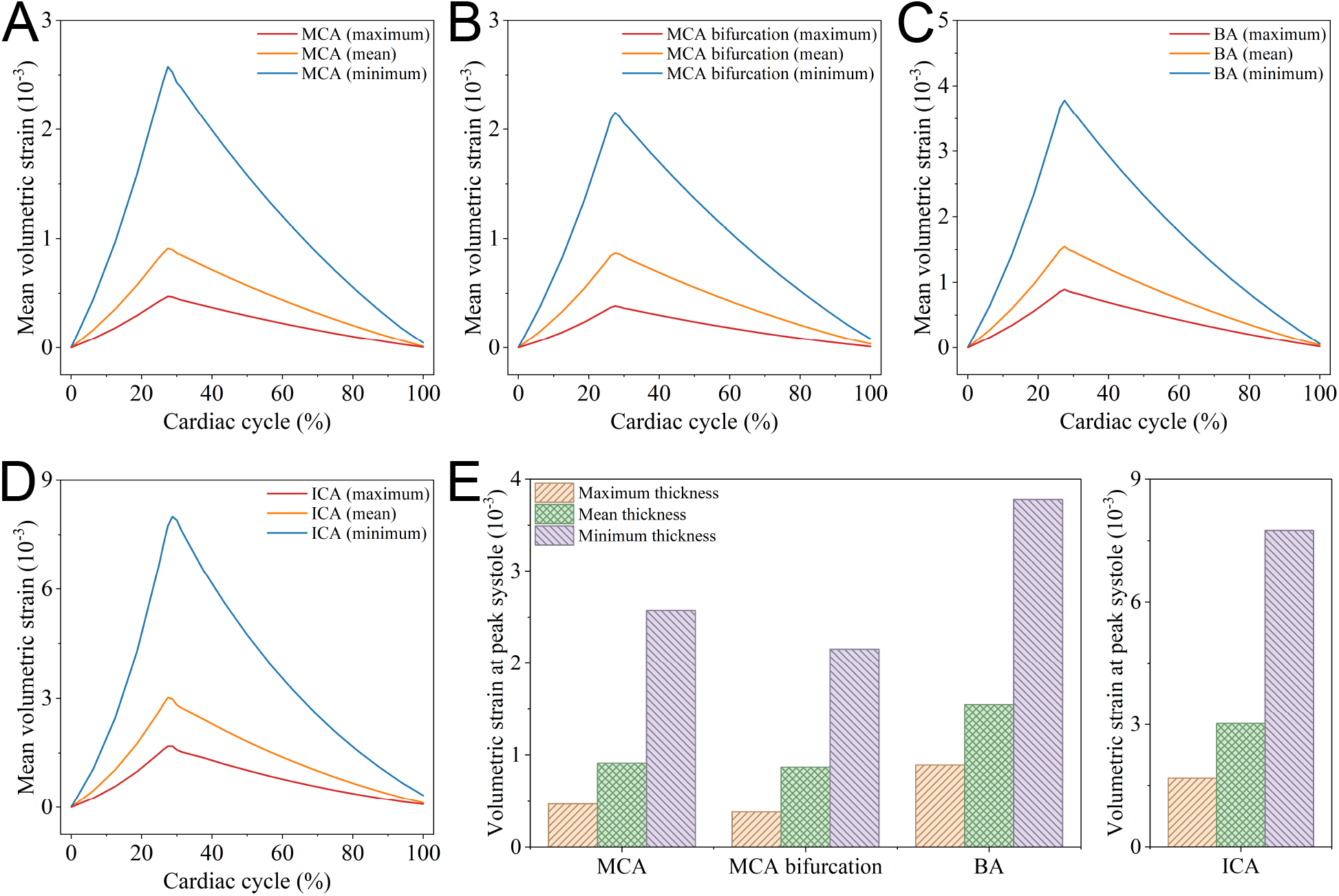
Effect of vessel wall thickness on volumetric strain. Mean volumetric strain throughout the cardiac cycle for the **(A)** MCA, **(B)** MCA bifurcation, **(C)** BA, and **(D)** ICA models using the maximum, mean, and minimum vessel wall thicknesses. **(E)** Corresponding volumetric strains at peak systole across four models. In all models, the minimum wall thickness results in the largest volumetric strain, while the maximum wall thickness produces the lowest strain. All volumetric strain data are extracted from the first tissue layer of each model.

As shown in Figure 7E, the peak-systolic results further demonstrate that the effect of vessel wall thickness is most pronounced in the ICA model, followed by the BA, MCA, and MCA bifurcation models. At peak systole, the ICA shows the highest sensitivity to thickness, with volumetric strain ranging from 1.68 *×* 10^−3^ to 7.75 *×* 10^−3^. In contrast, the MCA bifurcation exhibits the smallest variation, ranging from 0.38 *×* 10^−3^ to 2.15 *×* 10^−3^, while the MCA and BA models show intermediate responses. This can be attributed to the lower structural stiffness of thinner arterial walls, which allows greater deformation under the same pulsatile loading conditions. All volumetric strain data presented in this section are extracted from the first tissue layer of each model, as plotted in Figure 3A, which remains qualitative and is used for cross-comparison only.

### 3.4. Haemodynamic responses in the vicinity of the vessel wall

The haemodynamic responses near the vessel wall are evaluated using TAWSS, OSI and RRT, which are defined in Section 2.6. These parameters are analysed to assess the spatial variation of near-wall flow behaviour in the MCA, MCA bifurcation, BA and ICA models. In this section, all geometrical models used for evaluation are constructed using the mean vessel wall thickness.

Figure 8 shows the TAWSS distribution near the vessel wall. Overall, the distributions are heterogeneous among the different models. Relatively high TAWSS regions are observed along curved vessel segments, bifurcation regions and areas where the lumen diameter decreases. In the MCA model, TAWSS varies along the vessel surface, with locally elevated values appearing near the curved and distal regions. The MCA bifurcation model exhibits a more complex pattern, particularly around the branch region. In the BA model, the TAWSS distribution is comparatively smoother, although localised variations are still present near the vessel ends. For the ICA model, evident spatial variations in TAWSS are observed along the highly curved segments and near one of the narrowed ends, indicating the influence of vessel curvature and local geometrical changes on near-wall shear distribution.

**Figure 8:**
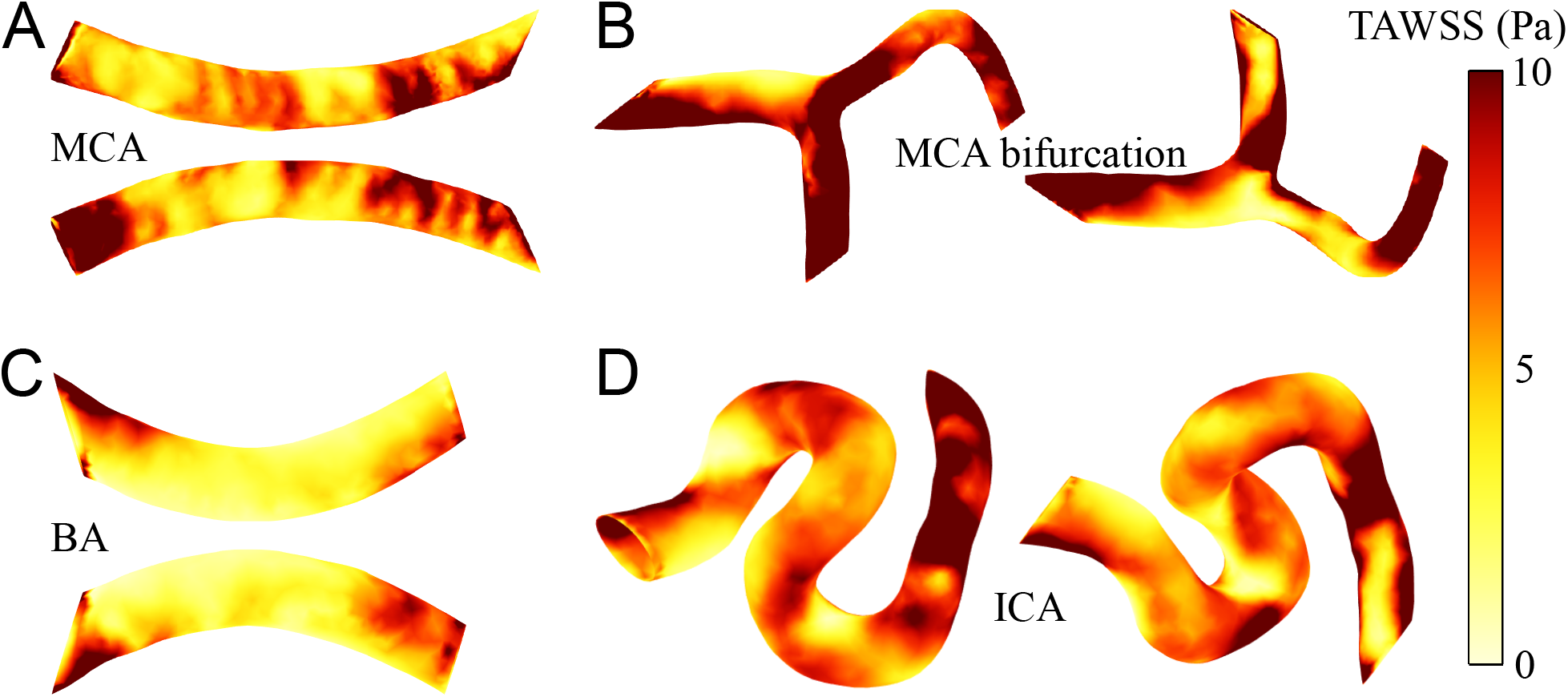
Surface contours of TAWSS distributions near the vessel wall in each vessel model, including **(A)** MCA, **(B)** MCA bifurcation, **(C)** BA, and **(D)** ICA. The colour scale represents TAWSS values ranging from 0 to 10 Pa.

Figure 9 and Figure 10 present the OSI and RRT distributions near the vessel wall, respectively. OSI values remain lower than 0.05 across most regions of the vessel surfaces in each model. Although spatial variations in low-magnitude OSI are visible on the logarithmic colour scale, the consistently low absolute values indicate that near-wall flow is predominantly unidirectional during the cardiac cycle. Similarly, RRT values are generally lower than 1 1/Pa in most vessel regions, suggesting limited near-wall flow residence. Nevertheless, localised elevations of both OSI and RRT are observed near geometrically complex regions, particularly at the MCA bifurcation, BA terminal regions and curved ICA segments. These regions are associated with disturbed near-wall flow, where curvature, branching, or local expansion promote flow separation, recirculation, and prolonged residence of blood particles near the vessel wall. The spatial correspondence between increased OSI and elevated RRT indicates that the behaviour of oscillatory flow contributes to a longer residence time in these localised regions.

**Figure 9:**
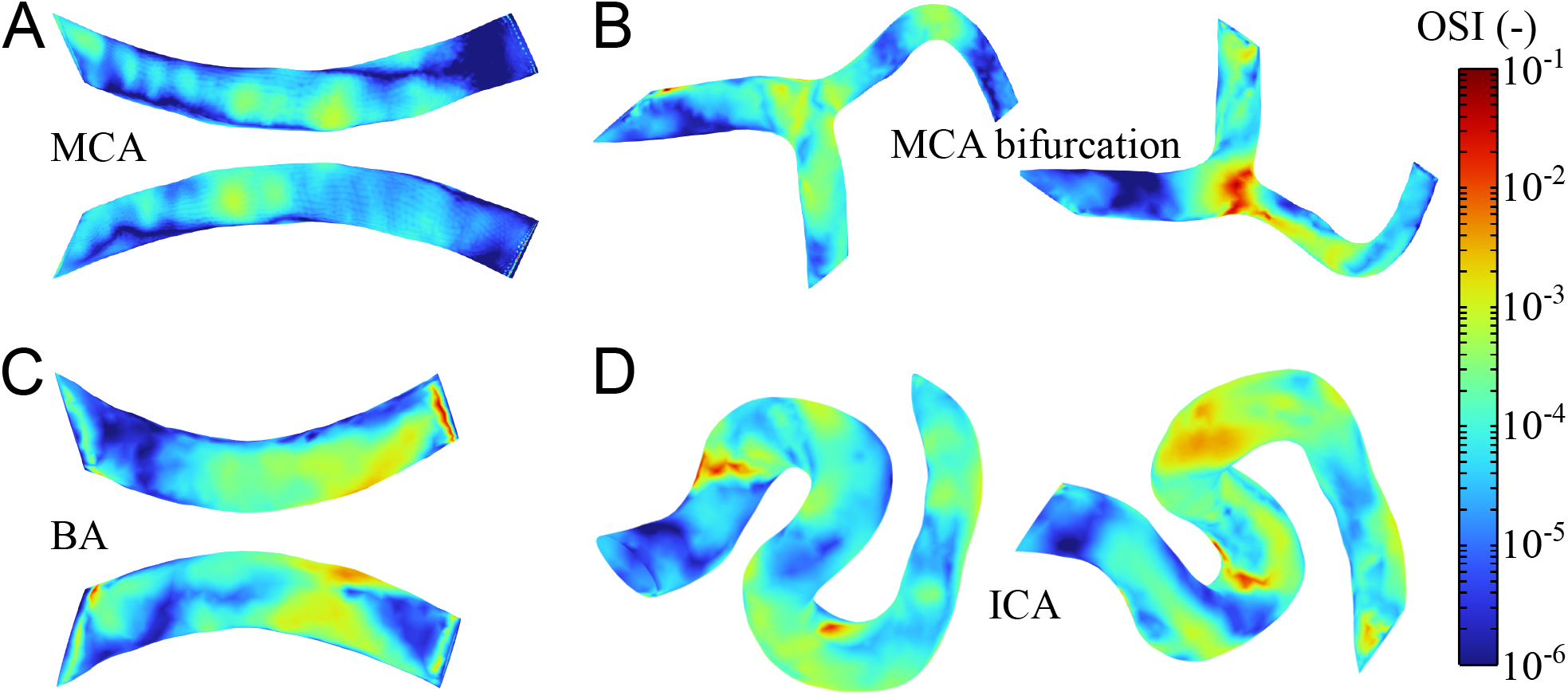
Surface contours of OSI distribution near the vessel wall in each vessel model, including **(A)** MCA, **(B)** MCA bifurcation, **(C)** BA, and **(D)** ICA. The logarithmic colour scale represents OSI values ranging from 10^−6^ to 10^−1^.

**Figure 10:**
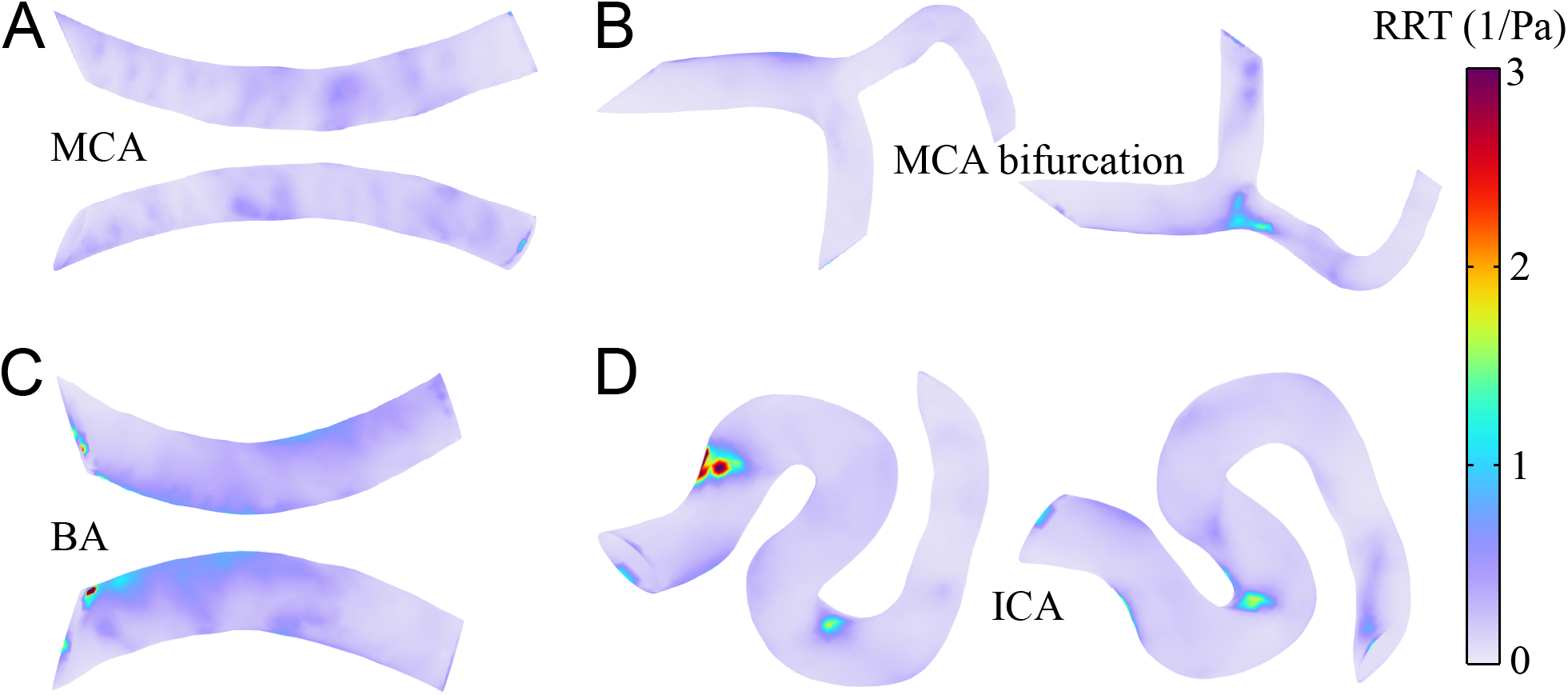
Surface contours of RRT distribution near the vessel wall in each vessel model, including **(A)** MCA, **(B)** MCA bifurcation, **(C)** BA, and **(D)** ICA. The colour scale represents RRT values ranging from 0 to 3 1/Pa.

To further compare the overall haemodynamic responses among the different vessel models, the surface-average WSS over a cardiac cycle, the surface-averaged TAWSS, OSI, RRT values are quantified, as shown in Figure 11. The surface-averaged WSS increased during the early stage of the cardiac cycle and reached its peak at approximately 30% of the cycle in all models, followed by a gradual decrease toward the end of the cycle (Figure 11A). Among the four models, the MCA bifurcation exhibits the highest surface-averaged WSS throughout the cardiac cycle, with a peak value of 12.56 Pa, whereas the BA model generally shows the lowest values. Although the MCA exhibits slightly higher surface-averaged WSS during most of the cardiac cycle, the ICA shows a marginally higher value during late diastole. This difference can be primarily attributed to the higher flow rate prescribed for the ICA, as well as the geometrical characteristics of the two vessels, which influence the redistribution of the near-wall velocity field during flow deceleration. Consistently, the surface-averaged TAWSS is highest in the MCA bifurcation model and lowest in the BA model (Figure 11B), with the values of 10.15 Pa and 5.11 Pa, respectively. For the surface-averaged OSI, the MCA bifurcation shows the largest value of 4.16 *×* 10^−4^, indicating a relatively higher degree of oscillatory near-wall flow compared with the other models (Figure 11C). In contrast, the BA model exhibits the highest surface-averaged RRT (Figure 11D), with a value of 0.24 1/Pa, suggesting that although its OSI is not the highest, the lower wall shear stress contributes to a longer relative residence time near the vessel wall.

**Figure 11:**
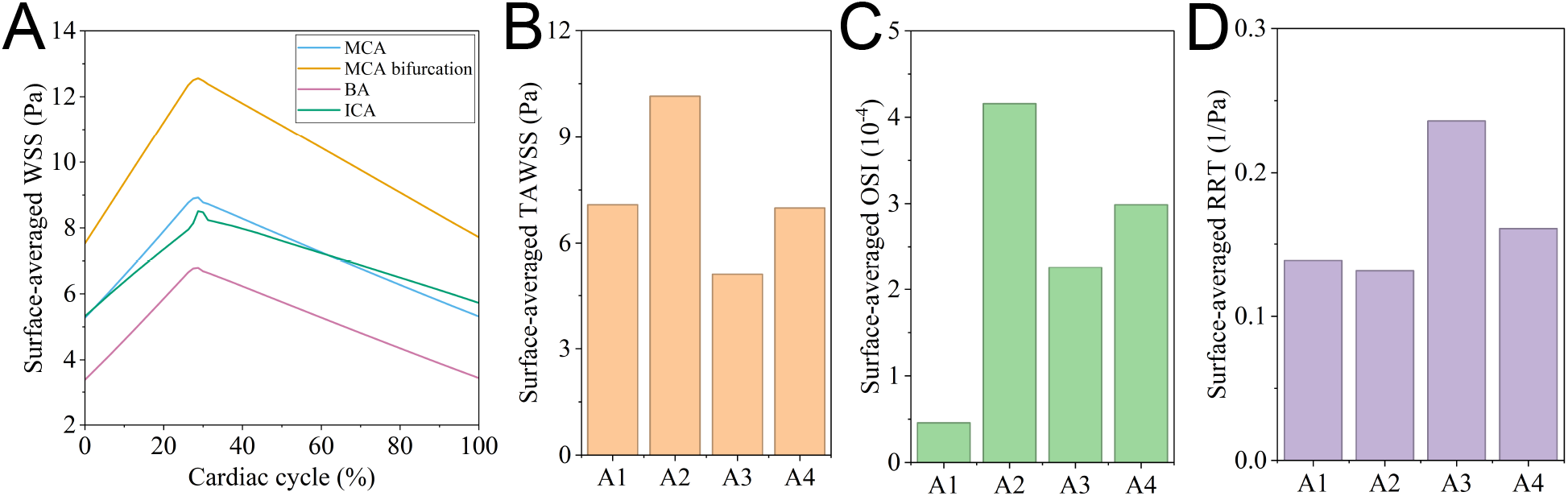
Quantitative comparison of surface-averaged haemodynamic parameters in different models. **(A)** Temporal variation of surface-averaged WSS during one cardiac cycle in each model. **(B)**-**(D)** Surface-averaged values of TAWSS, OSI and RRT, respectively. In panels **(B)**-**(D)**, A1, A2, A3 and A4 correspond to the MCA, MCA bifurcation, BA and ICA models, respectively.

## 4. Discussion

This study presents a coupled blood–vessel–brain dynamics modelling framework based on image-derived cerebral arterial geometries and validates its predictive accuracy against *in vivo* measurements of cardiac-induced brain deformation. The framework is subsequently used to investigate the coupled dynamic responses of blood flow, vessel motion, and surrounding brain tissue throughout the cardiac cycle. To the authors’ knowledge, this is the first model reported in the open literature to incorporate these three interacting components and to validate the resulting simulations against *in vivo* measurements. The findings indicate that local vascular geometry and vessel wall thickness substantially influence vessel-wall deformation, the associated mechanical response of the surrounding brain tissue, and near-wall haemodynamic behaviour.

The validation results indicate that the proposed mathematical framework can reasonably capture the experimentally measured cardiac-induced volumetric strain response. The simulated strain values generally follow the experimental temporal trend: increasing during systole and decreasing during diastole, and are mostly within the reported experimental range [17, 18]. In these referenced experimental studies, cerebral volumetric strain was measured using DENSE MRI, and peak volumetric strain showed good repeatability between repeated scans. The measurement also demonstrated spatial heterogeneity in the strain response, with grey matter exhibiting higher peak volumetric strain than white matter. In addtion, CSF and whole-brain tissue volume changes were strongly correlated over the cardiac cycle [17, 18]. This interaction is not explicitly represented in the present model, in which the CSF is excluded. Since the present validation compares simulated mean strain values with experimentally reported ranges rather than spatially matched voxel-wise measurements, it should be interpreted as an assessment of the overall magnitude and temporal behaviour of strain, rather than as a direct point-by-point reproduction of the experimental strain field. In addition, the experimental strain maps show localised regions with relatively high volumetric strain, which may be associated with vascular or perivascular pulsation effects [11, 13, 17, 18]. Since the present validation uses mean volumetric strain values, these local high-strain regions are averaged into the global tissue response and are not directly compared with the local strain concentrations predicted near the modelled arteries.

Differences between the simulated and experimental results may also reflect the distinct assumptions, formulations, and physical representations adopted in the two approaches. The experimental measurements [17, 18] were derived from image-based displacement fields and interpreted using an infinitesimal-strain formulation, whereas the computational model employs a finite-deformation framework with hyperelastic and hyper-viscoelastic constitutive descriptions of the vessel wall and surrounding brain tissue. Although the small-strain approximation is appropriate for the low-amplitude cardiac-induced deformations measured experimentally, differences in kinematic formulation, constitutive assumptions, vascular geometry, boundary conditions, and measurement resolution may contribute to discrepancies between the two sets of results. These discrepancies may be more pronounced in regions characterised by relatively large displacement gradients or substantial geometrical heterogeneity. In addition, cerebral arteries are not mechanically unconstrained *in vivo*. Their deformable walls are embedded within the intracranial environment and interact mechanically with the cerebrospinal fluid space, meninges, and surrounding brain parenchyma [6, 7, 8, 9]. These anatomical and mechanical interactions may restrict arterial motion and modify the local deformation and strain fields, but they are difficult to represent fully within a computational blood–vessel–brain model. Despite these limitations, the agreement between the simulated and experimentally measured strain ranges indicates that the proposed framework captures the principal characteristics of blood-flow-induced deformation of the cerebral vessel wall and adjacent brain tissue.

The temporal strain and displacement results show that pulsatile blood flow produces a clear mechanical response in both the vessel wall and the surrounding tissue. The strain and displacement responses are mainly concentrated near the vessel wall and gradually decreased away from the vascular structure, indicating that the influence of cardiac pulsation is primarily localised in the perivascular region. This finding suggests that cerebral arteries do not only serve as conduits for blood transport, but also act as dynamic mechanical sources that transmit cyclic loading to the surrounding brain tissue [1, 10, 13, 18]. Such pulsation-induced deformation may contribute to local tissue motion and perivascular mechanical stimulation [67, 44, 36, 68]. The magnitude and spatial distribution of this response are strongly affected by vessel geometry. Among the four investigated models, the ICA exhibits the largest deformation and displacement responses. This can be attributed to the combined effects of its higher blood flow rate and highly curved geometry, both of which can increase the mechanical loading acting on the vessel wall and its adjacent brain tissue. The higher flow rate produces greater haemodynamic forcing, while the pronounced curvature of the ICA alters the spatial distribution of pressure and flow-induced loading, resulting in a more heterogeneous deformation field [40]. Hence, the comparatively large response of the ICA should not be attributed to vessel curvature alone, but rather to the interaction between haemodynamic loading and vascular geometry. In comparison, the MCA, MCA bifurcation and BA models show smaller overall deformation magnitudes, although localised strain and displacement concentrations are still observed near curved regions, branch regions and vessel ends. Therefore, even when the global deformation response is relatively small, local geometric features can create regions of increased mechanical response. Since the four arterial models are extracted from a healthy subject and cover typical vascular features, including curvature, bifurcation and vessel-end regions, these results provide insight into representative blood-flow-induced deformation patterns in the healthy brain.

The analysis of vessel wall thickness further demonstrates that wall structure has a strong influence on the coupled mechanical behaviour. In all vessel models, thinner vessel walls produced larger volumetric strain, whereas thicker walls reduced the deformation response. This trend is mechanically reasonable because a thinner wall provides lower resistance to pulsatile pressure and flow-induced loading. The implication is that vessel wall thickness is not merely a geometric parameter, but a key determinant of how strongly vascular pulsation is transmitted to the surrounding tissue. Therefore, using a single uniform or mean wall thickness may be sufficient for general comparisons, but it would underestimate or overestimate local deformation in regions where wall thickness varies substantially [41]. This finding emphasises the importance of including the effect of wall thickness in future patient-specific or disease-specific simulations.

The haemodynamic results provide complementary insight into the near-wall flow environment and demonstrate that local vascular geometry strongly regulates wall-based flow indices. The heterogeneous TAWSS distributions indicate that wall shear exposure is not uniform across the vessel surface, but is concentrated in regions affected by curvature, bifurcations and local lumen variations [39, 40]. In contrast, OSI remains low over most vessel surfaces, indicating predominantly unidirectional near-wall flow during the cardiac cycle, while localised increases in OSI occur near geometrically complex regions. These localised OSI elevations are generally accompanied by increased RRT, suggesting that disturbed or oscillatory near-wall flow contributes to longer residence time near the vessel wall. However, the surface-averaged results in Figure 11 also indicate that RRT is not governed by OSI alone. For instance, the BA model exhibits a relatively high surface-averaged RRT despite not showing the highest OSI, which can be attributed to its lower wall shear stress level. This highlights that the RRT reflects the combined influence of low wall shear stress and oscillatory flow [65, 66]. A key implication of these results is that mechanical deformation and haemodynamic disturbance are both controlled by vascular geometry, but not necessarily in the same way. Regions with large deformation are not always identical to regions with high OSI or RRT. This indicates that a complete assessment of cerebral artery behaviour should consider both structural deformation indices and near-wall haemodynamic indices. Evaluating only one aspect would provide an incomplete description of the local blood–vessel–brain environment.

Beyond characterising cerebral haemodynamics and tissue mechanics, the present framework also provides a potential platform for investigating how cardiac pulsation influences transport and drug delivery within the brain. The cardiac-induced deformation quantified here produces cyclic changes in the local geometry and mechanical state of the tissue surrounding cerebral arteries, including regions through which interstitial and perivascular transport occurs. This is particularly relevant because our previous studies have demonstrated that intracerebral transport is highly sensitive to brain microstructure and its deformation. Pressure-induced changes in tissue microstructure can substantially alter porosity and hydraulic permeability [24, 69, 70], while axonal organisation and tissue anisotropy can strongly influence interstitial fluid flow, nanoparticle diffusion and the spatial distribution of therapeutic agents [71, 72, 73, 74]. More recently, microstructure-informed multiscale modelling has enabled the prediction of interstitial transport and drug-delivery pathways from the cellular to the whole-brain scale, further highlighting the importance of dynamically evolving tissue transport properties [35, 75, 76, 77]. The present blood–vessel–brain framework therefore provides a new opportunity to connect cardiac-induced vascular and tissue deformation with these transport mechanisms, enabling future studies to quantify how physiological arterial pulsation dynamically modulates interstitial and perivascular transport and, ultimately, the distribution of therapeutic agents within the brain.

Despite these insights in this work, several assumptions and simplifications should be considered and are worth further discussion. First, the surrounding brain tissue is represented by an idealised cuboid domain, within which the vessels are assumed to be fully embedded. This represents an additional geometric simplification, as *in vivo* vessels are not necessarily uniformly or completely surrounded by brain tissue, depending on their anatomical location and the surrounding structures [78]. Although this simplification does not fully capture the complex geometry, regional heterogeneity, and anatomical conditions of the *in vivo* brain environment [79, 80, 81], it provides the most appropriate modelling strategy for the present study. In particular, it enables controlled comparisons among the different vessel models while maintaining computational feasibility and limiting confounding effects arising from whole-brain anatomical complexity. With access to greater computational resources, future studies could extend the framework to anatomically realistic vessel-specific surrounding environments and whole-brain simulations with spatially varying tissue properties, thereby relaxing this limitation. Second, the vessel wall thickness is assumed to be uniformly distributed around the lumen, whereas cerebral arterial wall thickness can vary spatially along the vessel and around the circumference, particularly near curved regions, bifurcations and branch points [82, 83, 84, 85, 86]. This simplification may affect the predicted local deformation and near-wall haemodynamic indices. Third, the CSF flow and other anatomical constraints, including the meninges, perivascular spaces, etc., are not explicitly included in the present framework. These components could modulate the transmission of vascular pulsatility to the CSF and surrounding parenchyma by altering local compliance, hydraulic resistance, and mechanical boundary conditions, thereby affecting regional mechanical response of brain tissue [7, 87, 88, 89, 90]. However, the magnitude and relative contribution of these effects are not sufficiently quantified to parameterise them independently within the present model. Explicit inclusion would therefore introduce additional geometrical, mechanical, and fluid-dynamic parameters that are themselves poorly constrained. We consequently restrict the present framework to the blood-vessel-brain mechanical interaction in an idealised tissue domain, rather than to present the complete *in vivo* mechanical environment. Future work could extend the present framework by incorporating anatomically realistic brain geometries, spatially varying vessel-wall thicknesses, and patient-specific vascular and tissue characteristics, thereby improving the physiological fidelity of simulated blood–vessel–brain dynamics.

## 5. Conclusions

In this study, we develop an image-derived computational framework to analyse coupled blood-vessel-brain dynamics in representative cerebral arterial models. Four cerebral arterial segments are reconstructed from the high-field MRI data of a healthy subject and different thicknesses of the vessel wall are assigned to each model using their reported data. The simulated volumetric strain responses show reasonable agreement with reported experimental measurements, supporting the capability of the framework to capture the main mechanical features of cardiac-induced vessel and tissue deformation. The modelling results reveal that pulsatile blood flow induces localised deformation in both the arterial wall and surrounding brain tissue. The magnitude and spatial distribution of this response are strongly affected by vascular geometry, with the ICA model showing the largest deformation due to the combined effects of its higher blood flow rate and highly curved geometry. Vessel wall thickness is also found to play an important role, as thinner walls produced larger volumetric strain while thicker walls reduced the deformation response. The haemodynamic analysis show that near-wall flow behaviour is highly geometry-dependent. TAWSS is heterogeneous across the vessel surfaces, while localised increases in OSI and RRT are observed near curved, bifurcating and terminal regions. These results indicate that cerebral arterial geometry influences both mechanical deformation and near-wall haemodynamic conditions.

Overall, the present framework provides a grounded basis for studying image-derived blood-flow-induced vessel and brain tissue responses. As the present analysis are performed using healthy cerebral arterial geometries, the findings establish a reference framework for understanding normal pulsatile blood–vessel–brain interaction. This baseline provides a foundation for future disease-oriented investigations in which pathological changes in vascular geometry, wall thickness, material stiffness or surrounding tissue constraints can be evaluated by comparing with the normal response.

## CRediT authorship contribution statement

**Yi Yang**: Writing - original draft, Methodology, Software, Visualisation, Formal analysis. **Meiyao Wang**: Writing – review & editing, Resources, Visualisation, Formal analysis. **Yumin Liu**: Writing – review & editing, Resources, Formal analysis. **Wenbo Zhan**: Writing – review & editing, Methodology, Supervision, Formal analysis. **Daniele Dini**: Writing – review & editing, Methodology, Supervision, Funding acquisition, Formal analysis. **Tian Yuan**: Writing – review & editing, Methodology, Software, Funding acquisition, Formal analysis, Conceptualisation.

## Acknowledgements

Tian Yuan was supported by the Department for Science, Innovation and Technology (DSIT) and the Royal Academy of Engineering (RAEng) under the Research Fellowships scheme. Daniele Dini was supported by the DSIT, RAEng, and Shell via a Research Chair in Complex Engineering Interfaces (RCSRF2122-14-143). Daniele Dini and Yi Yang would like to acknowledge the support received from the EPSRC via the Grant EP/X033546/1. Yi Yang would like to acknowledge the support received from the Great Britain-China Educational Trust Award. Meiyao Wang would like to acknowledge the support received from Hubei Provincial Natural Science Foundation Program for Young Scientists (Category B) (2025AFB263) and Youth Project of the Science and Technology Innovation Cultivation Fund of Zhongnan Hospital of Wuhan University (CXPY2025133).

## Data availability

The datasets generated and/or analysed during the current study are available from the corresponding authors on reasonable request.

